# Transposable Elements are an evolutionary force shaping genomic plasticity in the parthenogenetic root-knot nematode *Meloidogyne incognita*

**DOI:** 10.1101/2020.04.30.069948

**Authors:** Djampa KL Kozlowski, Rahim Hassanaly-Goulamhoussen, Martine Da Rocha, Georgios D Koutsovoulos, Marc Bailly-Bechet, Etienne GJ Danchin

## Abstract

Despite reproducing without sexual recombination, the root-knot nematode *Meloidogyne incognita* is adaptive and versatile. Indeed, this species displays a global distribution, is able to parasitize a large range of plants and can overcome plant resistance in a few generations. The mechanisms underlying this adaptability without sex remain poorly known and only low variation at the single nucleotide polymorphism level have been observed so far across different geographical isolates with distinct ranges of compatible hosts. Hence, other mechanisms than the accumulation of point mutations are probably involved in the genomic dynamics and plasticity necessary for adaptability. Transposable elements (TEs), by their repetitive nature and mobility, can passively and actively impact the genome dynamics. This is particularly expected in polyploid hybrid genomes such as the one of *M. incognita*. Here, we have annotated the TE content of *M. incognita*, analyzed the statistical properties of this TE content, and used population genomics approach to estimate the mobility of these TEs across 12 geographical isolates, presenting phenotypic variations. The TE content is more abundant in DNA transposons and the distribution of TE copies identity to their consensuses sequence suggests they have been at least recently active. We have identified loci in the genome where the frequencies of presence of a TE showed variations across the different isolates. Compared to the *M. incognita* reference genome, we detected the insertion of some TEs either within genic regions or in the upstream regulatory regions. These predicted TEs insertions might thus have a functional impact. We validated by PCR the insertion of some of these TEs, confirming TE movements probably play a role in the genome plasticity with possible functional impacts.

## Introduction

Agricultural pests cause substantial yield loss to the worldwide life-sustaining production (Savary et al. 2019) and threaten the survival of different communities in developing countries. With a constantly growing human population, it becomes more and more crucial to reduce the loss caused by these pests while limiting the impact on the environment. In this context, understanding how pests evolve and adapt both to the control methods deployed against them and to a changing environment is essential. Among Metazoa, nematodes and insects are the most destructive agricultural pests. Nematodes alone are responsible for crop yield losses of ca. 11% which represents up to 100 billion € economic loss annually (Agrios 2005; McCarter 2009). The most problematic nematodes to worldwide agriculture belong to the genus Meloidogyne (Jones et al. 2013) and are commonly named root-knot nematodes (RKN) owing to the gall symptoms their infection leaves on the roots. Curiously, the RKN species showing the wider geographical distribution and infecting the broadest diversity of plants reproduce asexually via mitotic parthenogenesis (Trudgill and Blok 2001; Castagnone-Sereno and Danchin 2014). In the absence of sexual recombination, the genomes are supposed to irreversibly accumulate deleterious mutations, the efficiency of selection is reduced due to linkage between conflicting alleles while the combination of beneficial alleles from different individuals is impossible (Muller 1964; Hill and Robertson 1966; Kondrashov 1988; Glémin et al. 2019). For these reasons, asexual reproduction is considered an evolutionary dead end and is actually quite rare in animals (Rice 2002). In this perspective, the parasitic success of the parthenogenetic RKN might represent an evolutionary paradox.

Previous comparative genomics analyses have shown the genomes of the most devastating RKN are polyploid as a result of hybridization events (Blanc-Mathieu et al. 2017; Szitenberg et al. 2017). In the parthenogenetic RKN *M. incognita*, the gene copies resulting from allopolyploidy not only diverge at the nucleotide level but also in their expression patterns, suggesting this peculiar genome structure could support a diversity of functions and might be involved in the parasitic success despite the absence of sexual reproduction (Blanc-Mathieu et al. 2017). This hypothesis seems consistent with the ‘general purpose genotype’ concept, which proposes successful parthenogens have a generalist genotype with good fitness in a variety of environments (Vrijenhoek and Parker 2009). An alternative non mutually exclusive hypothesis is the ‘frozen niche variation’ concept which proposes parthenogens are successful in stable environments because they have a frozen genotype adapted to this specific environment (Vrijenhoek and Parker 2009). Interestingly, the frequency of parthenogenetic invertebrates is higher in agricultural pests, probably because the anthropized environments in which they live are more stable and uniform (Hoffmann et al. 2008).

However, although a general purpose genotype brought by hybridization might contribute to the wide host range and geographical distribution of these parthenogenetic RKNs, this alone, cannot explain how these species evolve and adapt to new hosts or environments without sex. For instance, initially, avirulent populations of some of these RKN, controlled by a resistance gene in a tomato, are able to overcome the plant resistance in a few generations, leading to virulent sub-populations, in controlled laboratory experiments (Castagnone-Sereno et al. 1994; Castagnone-Sereno 2006). Emergence of virulent populations, not controlled anymore by resistance genes have also been reported in the field (Barbary et al. 2015).

The mechanisms underlying the adaptability of parthenogenetic RKN without sex remain elusive. Recent population genomics analyses showed that only a few single nucleotide variations (SNV) could be identified by comparing different Brazilian *M. incognita* isolates showing distinct ranges of host compatibility (Koutsovoulos et al. 2020). Addition of further isolates from different geographical locations across the world did not substantially expand the number of variable positions in the genome. Furthermore, the few identified SNV showed no significant correlation with either the geographical location, the host range or the currently infected crop species. However, these SNV could be used as markers to confirm the absence of sexual meiotic recombination in *M. incognita*. Thus, the low nucleotide variability that was observed between isolates is probably not the main driver of the genomic plasticity underlying the adaptability of *M. incognita*.

Consistent with these views, convergent gene copy number variations were observed following resistance breaking down by two originally avirulent populations of *M. incognita* from distinct geographic origins (Castagnone-Sereno et al. 2019). The mechanisms supporting these gene copy numbers and other genomic variations possibly involved in the adaptive evolution of *M. incognita* remain to be described.

Transposable elements (TEs), by their repetitive and mobile nature, can both passively and actively impact genome plasticity. Being repetitive, they can be involved in illegitimate genomic rearrangements leading to loss of genomic portions or expansion of gene copy numbers. Being mobile, they can insert in coding or regulatory regions and have a functional impact on the gene expression or gene structure / function itself. For instance, TE neo-insertions have been shown to affect gene expression in a species-specific manner in amniotes (Zeng et al. 2018) and, in rodents, TE insertions account for ca. 20% of gene expression profile divergence between mice and rats (Pereira et al. 2009). At shorter evolutionary scales, differential presence / absence of TE across *Arabidopsis* populations revealed rare variants associated with extremes of gene expression (Stuart et al. 2016). TE insertions in coding regions can disrupt a gene and this disruption might eventually have an adaptive effect. For example, a TE insertion has caused disruption of a Phytochrome A gene in some soybean strains, which caused photoperiod insensitivity and was in turn associated with adaptation to high latitudes in Japan (Kanazawa et al. 2009). Moreover, in *Drosophila*, insertion of a TE in the *CHKov1* gene caused four new alternative transcripts and this modification is associated with resistance to insecticide and viral infection (Aminetzach et al. 2005; Magwire et al. 2011). In parallel, although TE movements can provide beneficial genomic novelty or plasticity, their uncontrolled activity can also be highly detrimental and put the organism at risk. For instance, some human diseases such as hemophilia (Kazazian et al. 1988) or cancers (Miki et al. 1992) are caused by TE insertions in coding or regulatory regions.

Concerning agricultural pests themselves, TEs are a major player of adaptive genome evolution by both passively and actively impacting the genome structure and sequence in some fungal phytopathogens (Faino et al. 2016). Whether TEs also play an important role in the genome plasticity and possibly adaptive evolution of parasitic animals, engaged in a continuous arms race with their hosts, remains poorly known. According to the Red Queen hypothesis, host-parasites arms race is a major justification for the prevalence of otherwise costly sexual reproduction (Lively 2010) and, in the absence of sex, other mechanisms should provide the necessary plasticity to sustain this arms race.

From an evolutionary point of view, the parthenogenetic root-knot nematode *M. incognita* represents an interesting model to study the activity of TEs and their impact on the genome, including in coding or regulatory regions. Indeed, being a plant parasite, *M. incognita* is engaged in an arms race with the plant defence systems and point mutations alone are not expected to be a major mechanism supporting adaptation in this species (Koutsovoulos et al. 2020).

In a broader perspective, little is known yet about the TE dynamics in nematode genomes and their possible impact on adaptive evolution, including in the model *C. elegans*, despite being the first sequenced animal genome since 1998 (The C. elegans Genome Sequencing Consortium 1998). Transposition activity of Tc1 TIR element was shown to be positively linked to the overall mutation rate in *C. elegans* mutator strains, one of which is characterized by high transposition in the germline, hence constituting a considerable evolutionary force (Bégin and Schoen 2007). However, these results may be hindered by the fact that, in wild-type *C. elegans*, although Tc1 excision frequency is substantial in somatic cells, it is negligible in the germ-cells (Emmons and Yesner 1984).

Besides Tc1, a more comprehensive analysis using population genomics approach in *C. elegans* represents the most advanced study of the TE dynamics in this species to date (Laricchia et al. 2017). By analyzing hundreds of wild populations of *C. elegans*, the authors have shown a substantial level of activity for multiple families of TEs in these genomes compared to the N2 reference strain. The study points at a population-wide variability of this activity, and, surprisingly, towards little evident phenotypic effect of this activity, even when TEs were found inserted into coding sequences. Concerning the possible functional impact of TE activity in nematodes, an investigation of TE expression in *C. elegans* germline in a single cell framework has shown significant differences between the expression pattern of LTR, non-LTR elements and DNA TE, associated with differentiated vs. undifferentiated cell types (Ansaloni et al. 2019). These complex cell-type specific differential expression patterns suggest TE activity plays an important role in the *C. elegans* embryonic development, although the exact role remains elusive. Overall, while it is now clearly established that TE are active in *C. elegans* and probably contribute to the genome plasticity, their possible functional implication or role in nematode adaptive evolution has not been shown so far.

In this study, we have tested whether the TE activity could represent a mechanism supporting genome plasticity in *M. incognita*, a prerequisite for adaptive evolution. We have re-annotated the 185Mb triploid genome of *M. incognita* (Blanc-Mathieu et al. 2017) for TEs, using stringent filters to identify canonical TEs, possibly active in the genome. We analyzed the statistical properties of the TE content and the distribution of TE sequence identity levels to their consensuses was used as a reporter of the recentness of their activity. We have then tested whether the frequencies of presence/absence of these TEs across the genome varied between different isolates. To test for variations in frequencies, we have used population genomics data from eleven *M. incognita* isolates collected on different crops and locations and differing in their ranges of compatible hosts (Koutsovoulos et al. 2020). From the set of TE loci that presented the most contrasted patterns of presence/absence across the isolates, we investigated whether some could represent neo-insertions. To estimate the possible functional impact of TE insertions, we checked whether some were inserted within coding or possible regulatory regions. Finally, we validated by PCR assays some of these neo-insertions in coding or regulatory regions, predicted by population genomics data. Overall, our study represents the first estimation of TE activity as a mechanism possibly involved in the genome plasticity and the associated functional impact in the most devastating nematode to worldwide agriculture. Besides *C. elegans*, little was known about the role of TE in the genome dynamics of *Nematoda*, one of the most species-rich animal phylum. Because this study focuses on an allopolyploid and parthenogenetic animal species, it also opens new evolutionary perspectives on the fate and potential adaptive impact of TEs in these singular organisms.

## Results

### The *M. incognita* TE landscape is diversified but mostly composed of DNA transposons

We used the REPET pipeline (Quesneville et al. 2005; Flutre et al. 2011) to predict and annotate the *M. incognita* repeatome (see methods). Here, we define the repeatome as all the repeated sequences in the genome, excluding Simple Sequence Repeats (SSR or microsatellites). The repeatome spans 26.38 % of the *M. incognita* genome length (sup.Table S1). As we wanted to assess whether TEs actively contributed to genomic plasticity, we applied a series of stringent filters on the whole repeatome to retain only repetitive elements presenting canonical signatures of TEs (see methods and (Kozlowski 2020a)). We identified 480 different TE-consensus sequences that allowed annotation of 9,633 canonical TE, spanning 4.67% of the genome (Table 1). Both retro (Class I) and DNA (Class II) transposons (Wicker et al. 2007) compose the *M. incognita* TE landscape with 5/7 and 4/5 of the known TE orders represented respectively, showing a great diversity of elements (Fig 1). Canonical retro-transposons and DNA-transposons respectively cover 0.90 and 3.77 % of the genome. Terminal Inverted Repeats (TIR) and Miniature Inverted repeat Transposable Elements (MITEs) DNA-transposons alone represent almost two-thirds of the *M. incognita* canonical TE content (64.49 %). Hence, the *M. incognita* TE landscape is diversified but mostly composed of DNA-transposons.

**Table 1:** Per-order summary of *M. incognita* canonical TE annotations. Autonomous TE orders (*) regroup elements known to present transposition machinery and thus able to transpose by themselves. On the opposite, non-autonomous orders (**) regroup elements lacking transposition machinery and therefore relying on autonomous elements to transpose.

|  | order<br>autonomous (*)<br>/<br>non-autonomous (**) | nb. of<br>features | total length<br>(bp) | genome<br>percentage<br>(%) | median<br>length (bp) | median identity<br>with<br>consensus (%) |
| --- | --- | --- | --- | --- | --- | --- |
| **Retro**<br>**-**<br>**transposon** | SINE (**) | 9 | 4,522 | 0.002 | 528.0 | 99.7 |
|  | LARD (**) | 45 | 6,342 | 0.035 | 1433.0 | 97.05 |
|  | TRIM (**) | 174 | 104,018 | 0.057 | 525.0 | 97.7 |
|  | LINE (*) | 145 | 313,224 | 0.171 | 1971.0 | 96.6 |
|  | LTR (*) | 373 | 1,164,836 | 0.635 | 2415.0 | 97.0 |
| **DNA**<br>**-**<br>**transposon** | Helitron (*) | 18 | 86,666 | 0.047 | 5080.0 | 94.4 |
|  | Maverick (*) | 189 | 1,307,068 | 0.712 | 6224.0 | 95.3 |
|  | MITE (**) | 5085 | 2,755,381 | 1.501 | 525.0 | 96.2 |
|  | TIR (*) | 3595 | 2,777,270 | 1.513 | 737.0 | 97.3 |
|  | **Total** | **9,633** | **8,576,405** | **4.673** |  |  |

**Fig 1:**
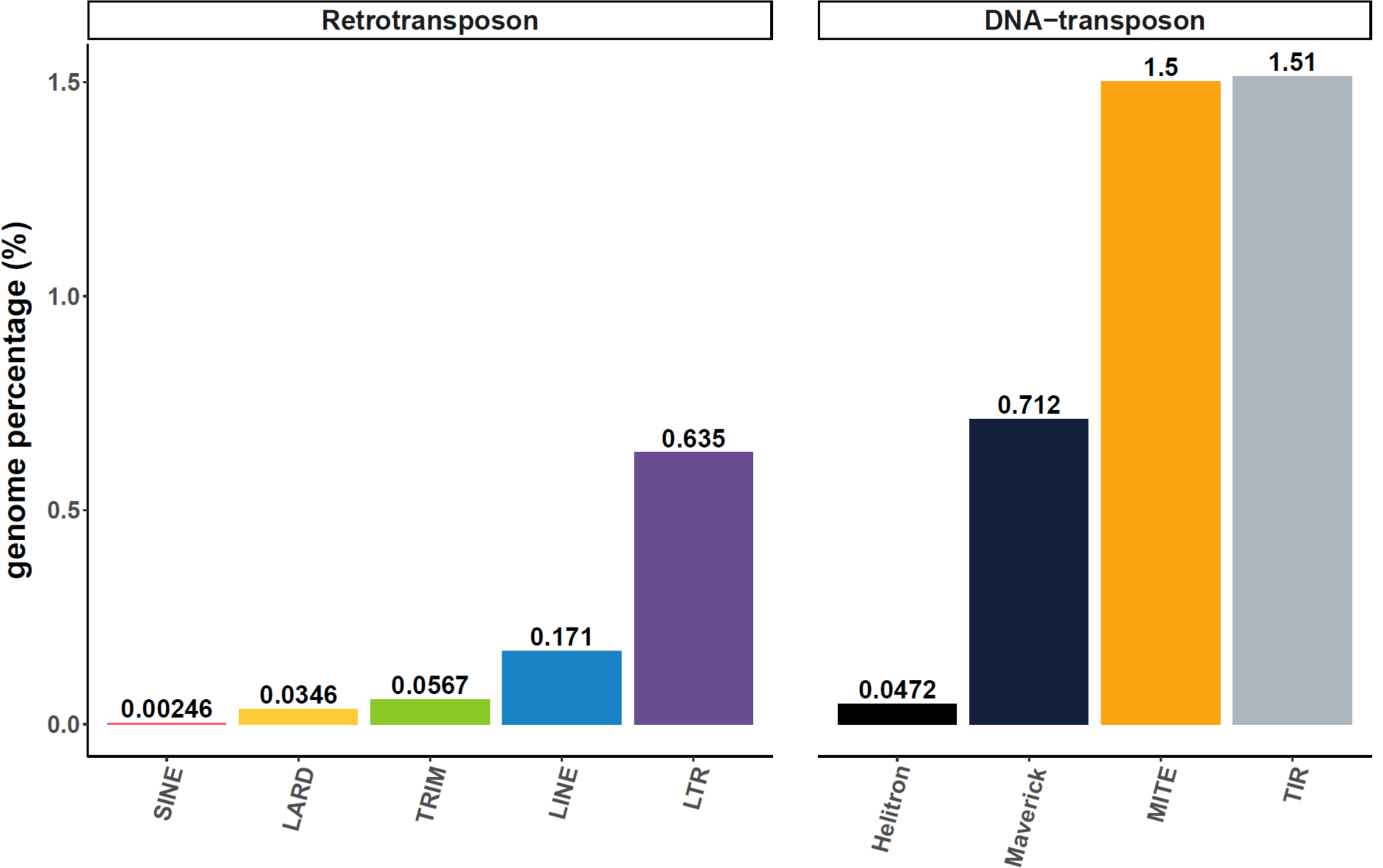
Canonical TE annotations distribution in *M. incognita* genome. Genome percentage is based on a *M. incognita* genome size of 183,531,997 bp (Blanc-Mathieu et al. 2017).

As a technical validation of our repeatome annotation protocol (see methods; sup. Fig S7), we performed the same analysis in *C. elegans*, using the PRJNA13758 assembly (The C. elegans Genome Sequencing Consortium 1998). We compared our results (Kozlowski 2020b) to the reference report of the TE landscape in this model nematode (Bessereau 2006) (sup. Table S2). We estimated that the *C. elegans* repeatome spans 11.81% of its genome, which is close to the 12 % described in (Bessereau 2006). The same resource also reported that MITEs and LTR respectively compose ∼2% and 0.4% of the *C. elegans* genomes while we predicted 1.8% and 0.2%. Predictions obtained using our protocol are thus in the range of previous predictions for *C. elegans;* which suggest our repeatome prediction and annotation protocol is accurate.

The wormbook resource (Bessereau 2006) mentioned that most of *C. elegans* TE sequences “are fossil remnants that are no longer mobile”, and that active TEs are DNA transposons. This suggests a stringent filtering process is necessary to isolate TEs that are the most likely to be active (e.g. the ‘canonical’ ones). Using the same post-processing protocol as for *M. incognita*, we estimated that canonical TEs span 3.60% of the *C. elegans* genome, with DNA-transposon alone representing 76.6% of these annotations (sup. Fig S1 & sup Table S3).

### Canonical TE annotations are highly identical to their consensus sequences and some present evidence for transposition machinery

Canonical TE annotations have a median nucleotide identity of 97% with their respective consensus sequences, but the distribution of identity values varies between TE orders (Fig 2, sup. Table S4). Most of the TEs within an order share a high identity level with their consensuses, the lowest values being observed for Helitron and Maverick elements. Yet, more than half of those elements share above 94% identity with their consensuses, (sup. Fig S3). Although it might be hypothesized the lower identities would be due to bigger length, we showed no evident correlation between the % identity copies share with their consensus and the proportion of consensus length covered (sup. Fig S3). Even considering our inclusion threshold at minimum 85% identity (see methods), the overall distribution of average % identities tends to be asymmetrical, and skewed towards higher values (Fig 2).

**Fig 2:**
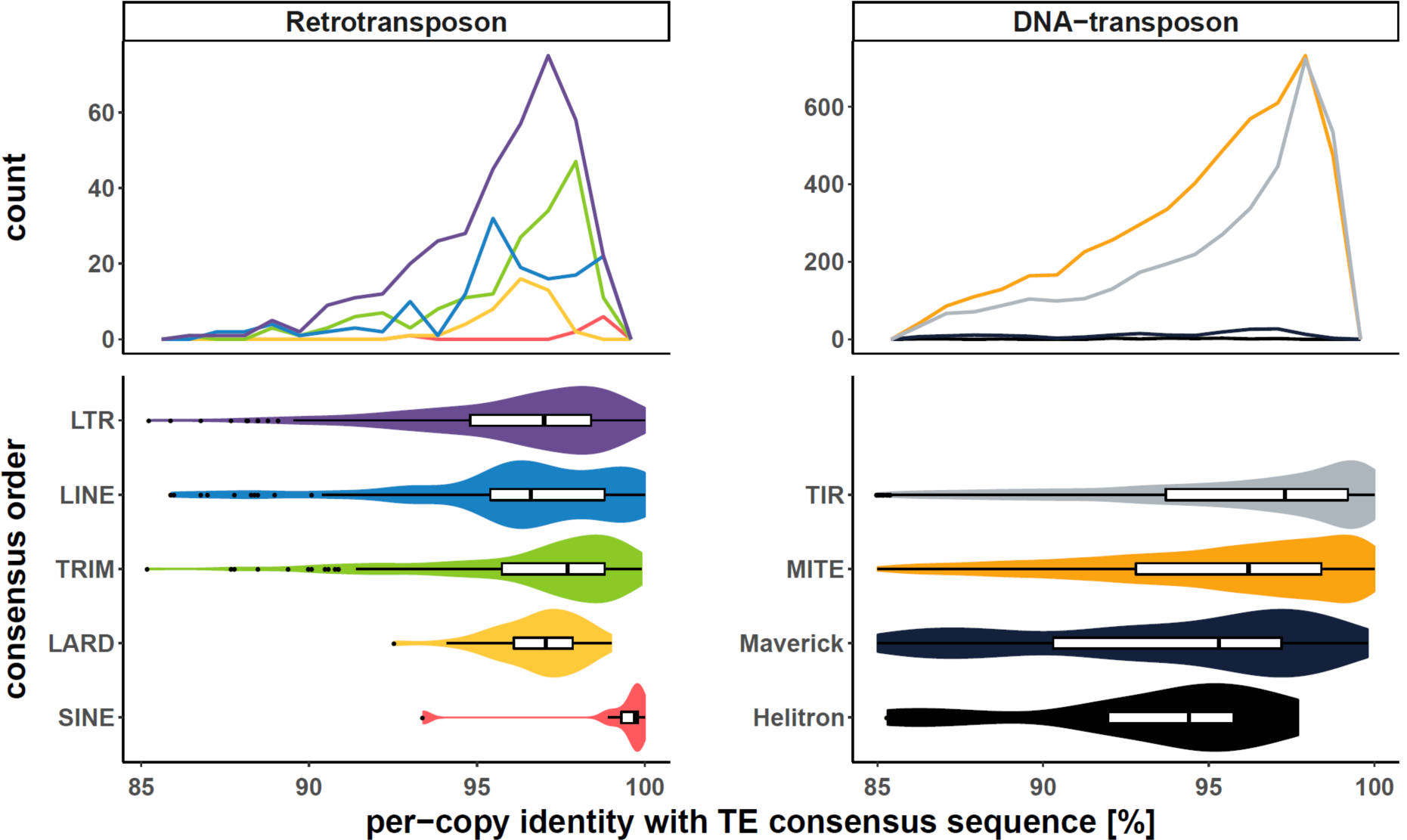
per-copy identity rate with consensus. Top frequency plots show the distribution of TE copies count per order in function of the identity % they share with their consensus sequence. To facilitate inter-orders comparison, bottom violin plots display the same information as a density curve, but also encompass boxplots. Each colour is specific to a TE order.

Among DNA-transposons, identity profiles of MITEs and TIRs to their consensuses were the most shifted to high values; one fourth of the TIRs annotations sharing above 99% identity with their consensus (Fig 2; sup. Tables 2 and 4).

Among retrotransposon, SINEs (present in very low numbers) and TRIMs show similar profiles with a quite narrow peak at more than 97% identity. Overall, these results indicate that notwithstanding small differences between orders, the canonical TEs show a high similarity with their consensuses.

High identity of TE annotations to their consensus can be considered a proxy of their recent activity (Bast et al. 2015; Lerat et al. 2019). To further investigate whether some TEs might be (or have been recently) active, we searched for the presence of genes involved in the transposition machinery within *M. incognita* canonical TEs (see methods). Among the canonical TE annotations, 6.21% (598/9,633) contain at least one predicted protein-coding gene, with a total of 893 genes involved. Of these 893 genes, 344 code for proteins with at least one conserved domain known to be related to transposition machinery. We found that 31.98% (110/344) of the transposition machinery genes had substantial expression support from RNA-seq data. In total, 106 canonical TE-annotations contain at least one substantially expressed transposition machinery gene (Kozlowski, Da Rocha, et al. 2020). These 106 TE annotations correspond to 39 different TE-consensuses, and as expected, only consensuses from the autonomous TE orders, e.g. LTRs, LINEs, TIRs, Helitron, and Maverick present TE-copies with substantially expressed genes coding for transposition machinery (sup. Table S5). Conversely, the non-autonomous TEs do not contain any transposition machinery gene at all. This suggests that some of the detected TEs have functional transposition machinery, which in turn could be hijacked by the non-autonomous elements.

Overall, the presence of a substantial proportion of TE annotations highly similar to their consensuses combined with the presence of genes coding for the transposition machinery and supported by expression data suggest some TE might be active in the genome of *M. incognita*.

### Thousands of loci show variations in TE presence frequencies across *M. incognita* isolates

We used the PopoolationTE2 (Kofler et al. 2016) pipeline on the *M. incognita* reference genome (Blanc-Mathieu et al. 2017) and the canonical TE annotation to detect variations in TE frequencies across the genome between 12 geographical isolates (see methods; (Kozlowski 2020b); sup. Fig S7). One isolate comes from Morelos in Mexico, which is the isolate that was used to produce the *M. incognita* reference genome. The 11 other isolates come from different locations across Brazil, and present four different ranges of compatible hosts (referred to as R1, R2, R3, R4, see sup. Fig S4) and currently infected crop species (Koutsovoulos et al. 2020). Pool-seq paired-end Illumina data has been generated for all these isolates. For each locus, each isolate has an associated frequency value representing the proportion of individuals in the pool having the TE detected at this location.

We identified 3,514 loci where the frequency variation between at least two isolates was higher than our estimated PopoolationTE2 error rate (0.00972 *i*.*e*. less than 1%, see methods).

Overall, the distribution of within-isolate frequencies is bimodal (Fig 3-A), and this pattern is common to all the isolates, including the reference Morelos isolate (Fig 3-B). On average, 21.1% of the loci have within-isolate frequencies < 25%, 60.7% have frequencies > 75%, and only 18.2% show intermediate frequencies Hence, most of the within-isolate TE frequencies pack around extreme values e.g. <25% or >75%.

**Fig. 3:**
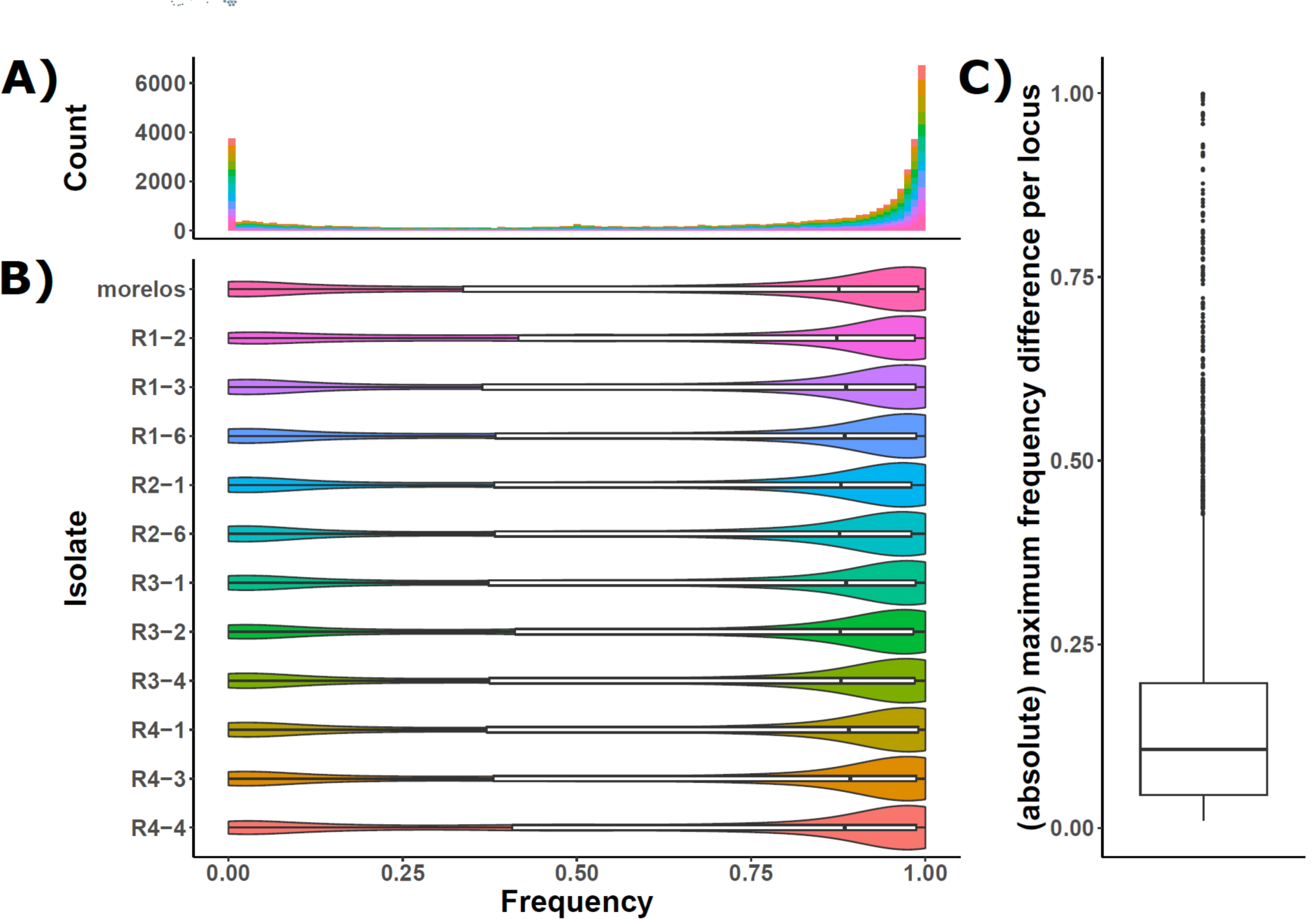
TE frequency distribution. The histogram (A) and violin plot (B) represent the TE frequency distribution per isolate. The colour chart is identical between the two figures. Both representations reveal that in all the isolates, only a few TE are found with intermediate frequencies. Right boxplot (C) represents the frequency absolute maximum difference per locus. For a given locus, it illustrates the frequency variability between isolates. The higher is the value; the more important is the frequency difference between at least two isolates. A value of 1 implies that the TE is absent in at least one isolate while it is present in 100% of the individuals of at least another isolate.

**Fig 4:**
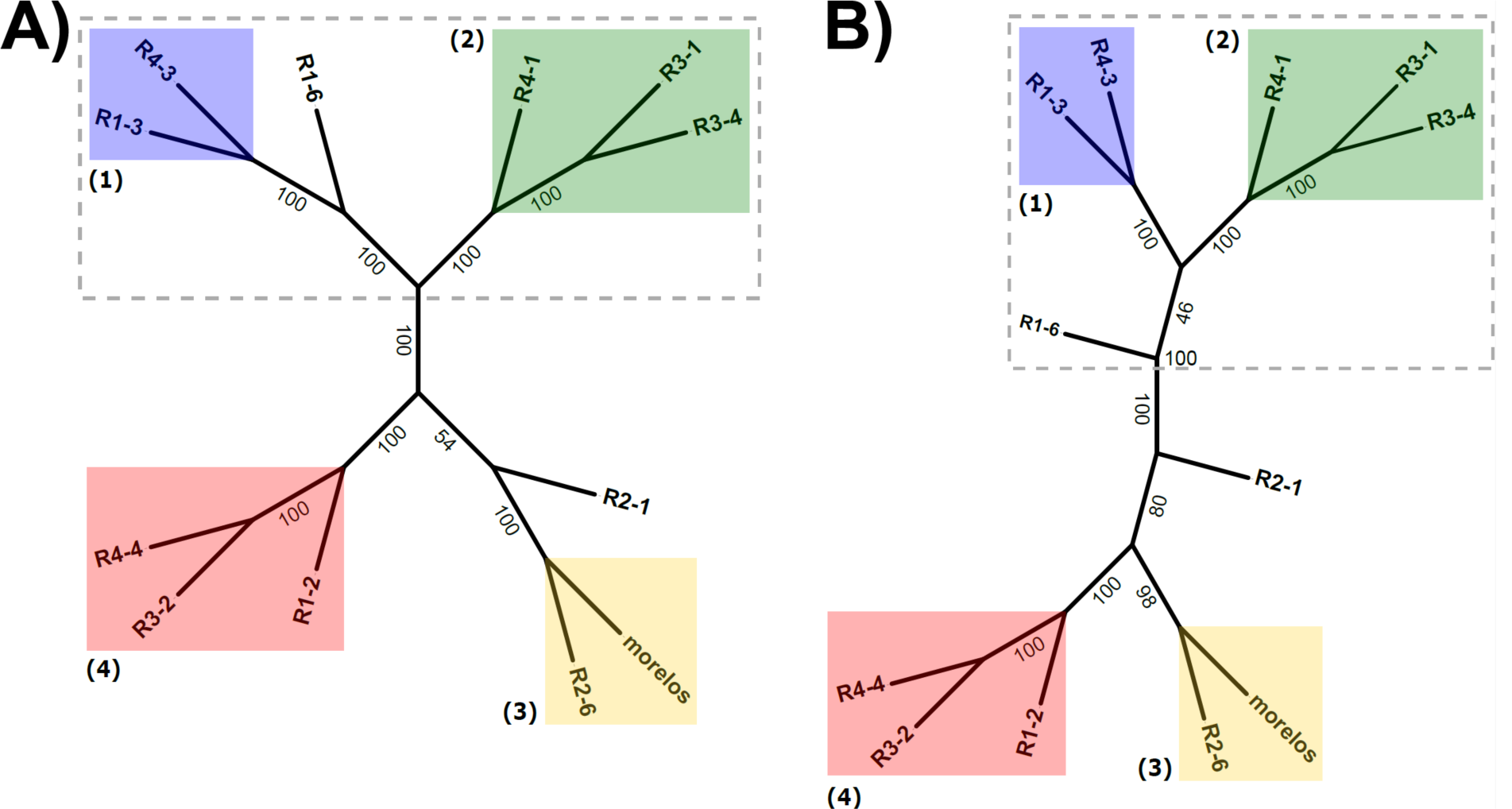
Phylogenetic tree for *M. incognita* isolates. A-Phylogenetic tree based on SNV present in coding sequences. Maximum Likelihood (ML) tree reconstruction. Branch length not displayed (see sup. Fig S5 for a version with branch length displayed). B-Phylogenetic tree based on TE-frequencies euclidean distances between isolates. Neighbor-Joining (NJ) tree reconstruction. Branch length not displayed (see sup. Fig S5 for a version with branch length displayed). In both trees, bootstrap support values are indicated on the branches. Isolates enclosed in the dashed area form a super-cluster composed of the clusters (1) and (2), and the isolate R1-6.

Nevertheless, these statistics provide no information about the frequency variability between isolates for a given locus. To address this question, for each locus, we computed the absolute maximum frequency difference between isolates (Fig 3-C). We found that the maximum frequency variation across the isolates is smaller than 20% in 75% of the loci (2,634/3,514). Hence, most of the loci show little to moderate isolate-wide variations in frequencies. Combined to the previous result, this implies that for most loci, the TEs are present either at a high or a low frequency among all isolates. However, some TE loci show more contrasted variations and will be the focus of further studies in our pipeline.

### Variations of TE frequencies across isolates recapitulate their divergence at the sequence level

We performed a Neighbour-joining phylogenetic analysis of *M. incognita* isolates based on a distance matrix constructed from TE frequencies (3,514 loci; see methods). We also performed a Maximum Likelihood (ML) analysis based on SNV in coding regions as previously identified in (Koutsovoulos et al. 2020) adding the reference isolate Morelos.

As shown in Fig.4, the TE-based and SNV-based tree topologies are highly similar. In particular, the two trees allowed defining four highly supported clades, with bootstrap support values ≥ 98. The four clades were identical, including branching orders for clades 2 and 4 (the two other clades containing each only two isolates). R1-6 and R2-1 positions slightly differed between the SNV-based (A) and TE-based (B) trees. However, in both trees, R1-6 is more closely related to clusters 1 and 2 than the rest of the isolates, and similar observations can be drawn for R2-1 with clusters 3 and 4.

Altogether, the similarity between the SNV-based and TE frequency-based trees indicates that most of the phylogenetic signal coming from variations in TE-frequencies between isolates recapitulates the SNV-based genomic divergence between isolates.

### Most of the TE frequency variations across the isolates concern TE present in the reference genome although additional TE loci were identified

As explained below (see also methods sup. Figs S7 & S8), we categorized all the loci with TE frequency variations between the isolates by (i) comparing their position to the TE annotation in the reference genome, (ii) analysing TE frequency in the reference isolate Morelos, (iii) comparing TE-frequencies detected for each isolate to the reference isolate Morelos. This allowed defining, on the one hand, non-polymorphic and hence stable reference annotation, and on the other hand, 3 categories of polymorphic (variable) loci (Fig 5).

**Fig 5:**
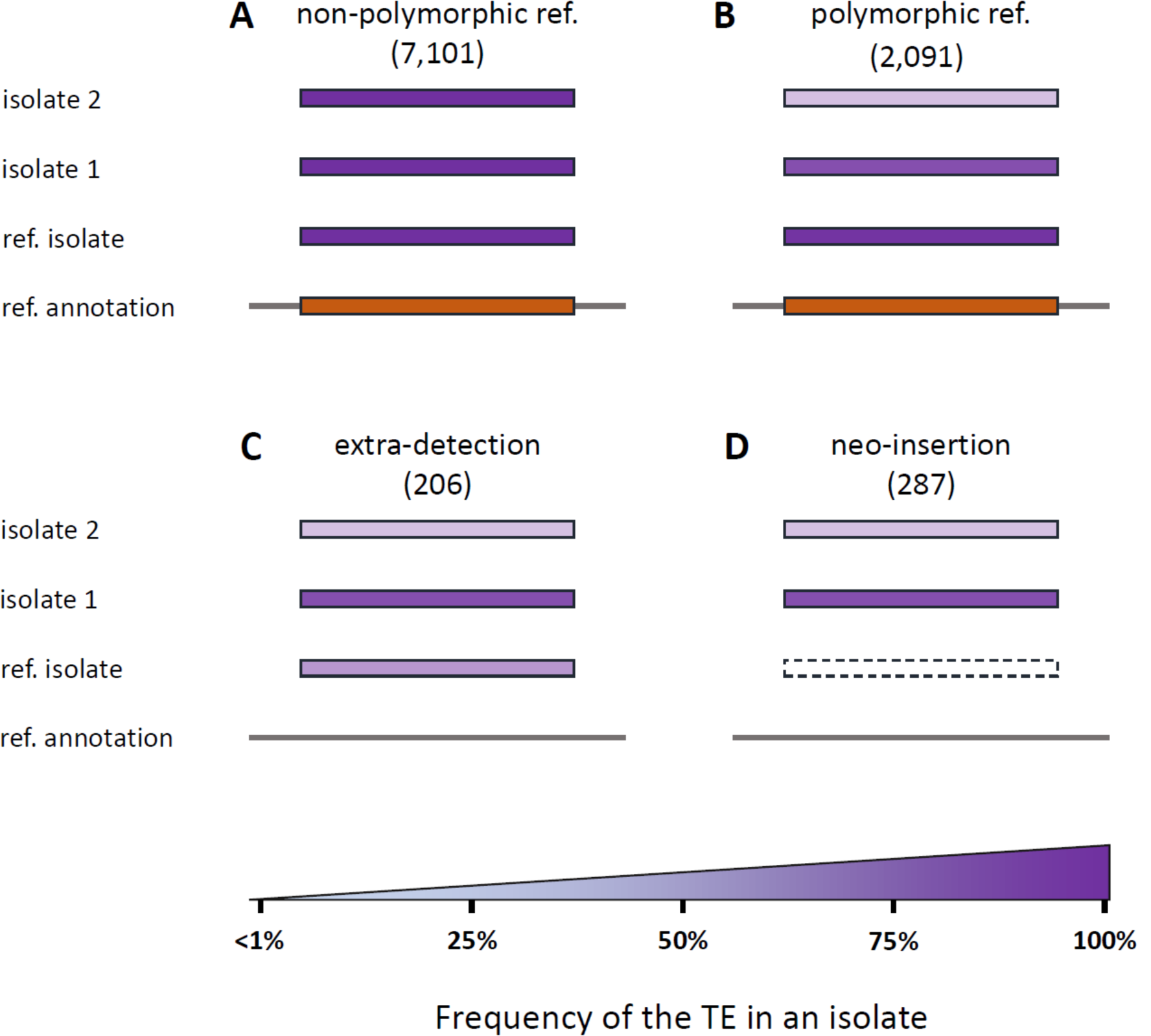
Categories of polymorphic TE loci. Orange boxes illustrate the presence of a TE at this locus in the reference genome annotation. Purple boxes illustrate the percentage of individuals in the isolates for which the TE is present at this locus (i.e. frequency). Frequency values are reported as colour gradients. A - non-polymorphic ref. TE locus: a TE is predicted in the reference annotation (orange box) AND no frequency variation exceeding 1% between isolates (Morelos included) is detected. B - polymorphic ref. locus: a TE is predicted in the reference annotation, is detected in the reference isolate Morelos with a frequency > 75%, and the presence frequency varies (>1%) in at least one isolate. C - extra-detection: no TE is predicted at this locus in the reference annotation but one is detected at a frequency >25% in the reference isolate Morelos, and optionally in other isolates. D - neo-insertion: no TE is predicted at this locus in the reference genome annotation and none is detected in the reference isolate (dashed box, frequency < 1%), but a TE is detected in at least another isolate with a frequency >= 25%.

Overall, 73.5% (2,584/3,514) of the loci with TE frequency variations could be assigned to one of the 3 categories of TE-polymorphisms (B, C, D in Fig 5) and the decomposition per TE order is given in Fig 6 and sup. Table S6.

**Fig 6:**
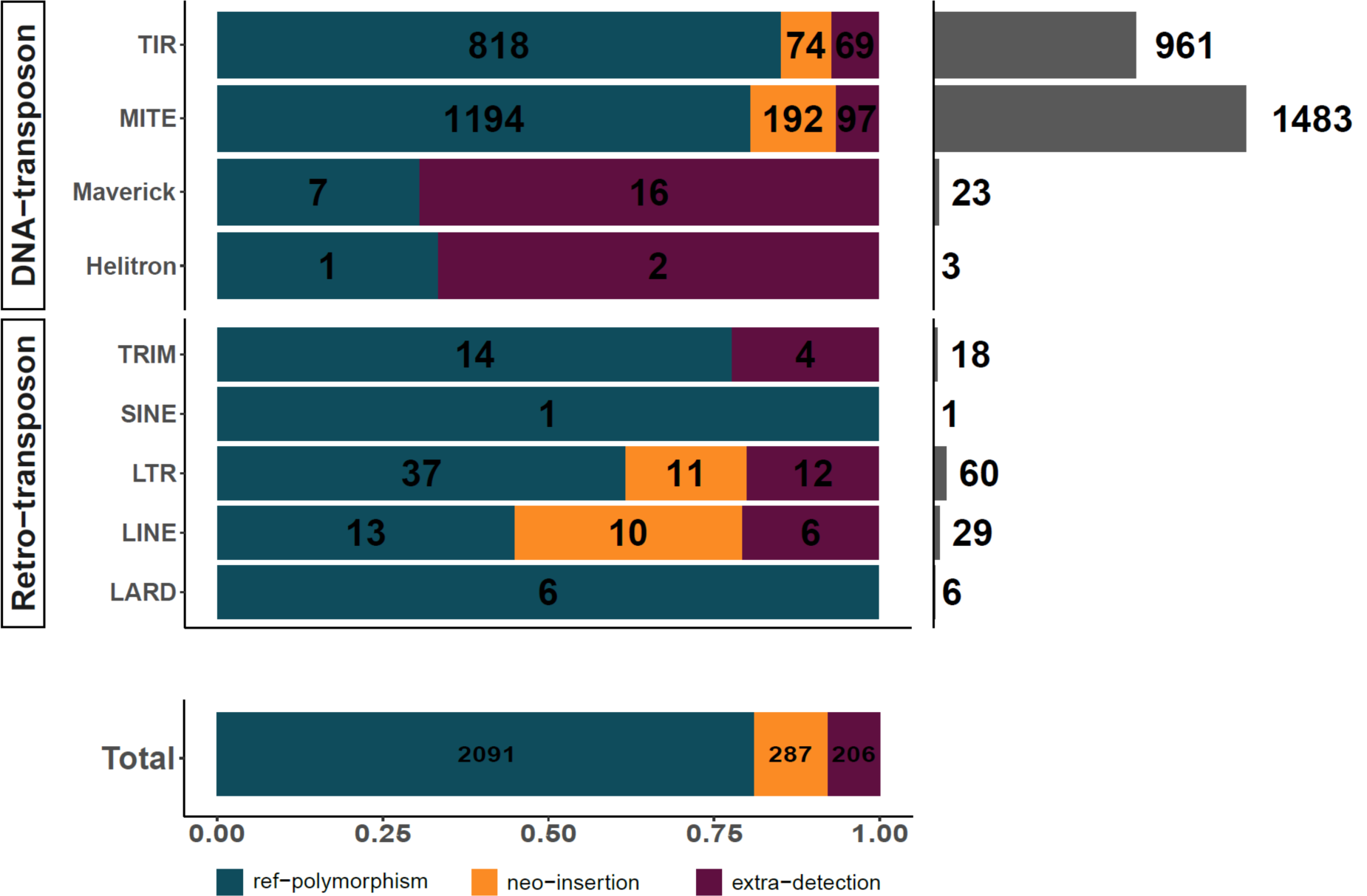
TE polymorphisms count per orders and types. The top left barplot shows TE polymorphisms distribution per type and per order. The bottom-left barplot summarizes TE polymorphisms distribution per type. In both barplots, the values in black represent the count per polymorphism type. The top-right barplot illustrates the total number of polymorphisms per order.

The vast majority of the polymorphic loci (80.92 %; 2,091/2,584) corresponds to an already existing TE-annotation in the reference genome and the corresponding TE is fixed (frequency > 75 %) at least in the reference isolate Morelos but varies in at least another isolate. These polymorphic loci cover ∼21.6% (2,091/9,702) of the canonical TE annotations, in total. These loci will be referred to as ‘polymorphic reference loci’ from now on (Fig 5B) and they encompass both DNA- and Retro-transposons.

Then, we considered as ‘neo-insertion’ TEs present at a frequency >25% in at least one isolate at a locus where no TE was annotated in the reference genome and the frequency of TE presence was higher than the estimated error rate (∼1%) in the reference Morelos isolate (Fig 5D). In total, 11.11 % (287/2,584) of the detected TE polymorphisms correspond to such neo-insertions. It should be noted here that we consider neo-insertions as regard to the reference Morelos isolate only and some of these so-called neo-insertions might represent TE loss in Morelos. Comparison with the phylogenetic pattern of presence / absence will allow distinguishing further the most parsimonious of these two possibilities (see next sections).

Finally, we classified as ‘extra-detection’ (Fig 5C) (7.97%; 206/2,584) the loci where no TE was initially annotated by REPET in the reference genome, but a TE was detected at a frequency >25% at least in the ref isolate Morelos by PopoolationTE2. It should be noted that 58.73% (121/206) of these loci correspond to draft annotations that have been discarded during the filtering process to only select the canonical annotations. These draft annotations might represent truncated or diverged versions of TE that exist in a more canonical version in another locus in the genome. Half of the remaining ‘extra-detections’ (42/85) are detected with low to moderate frequency (<42.6%) in the reference isolate Morelos. We hypothesise that because they represent the minority form, these regions were not taken into account during the assembly of the genome. This would explain why these TEs could not be detected in the genome assembly by REPET (assembly-based approach) but were identified with a read mapping approach on the genome plus repeatome by PopoolationTE2. The remaining ‘extra-detections’ might correspond to REPET false negatives, PopoolationTE false positives, or a combination of the two. Nonetheless, we can notice these cases only represent 1.63% (42/2,584) of the detected polymorphic TEs.

### TIR and MITE elements are overrepresented among TE-polymorphisms

By themselves, MITE and TIR elements encompass 94.58% (2,444/2,584) of the categorized TE-polymorphisms (Fig 6).

We showed that the polymorphism distribution varies significantly between the four categories presented in Fig. 5 (Chi-square test, p-value < 2.2e-16), indicating that some TE orders are characterised by specific polymorphisms types.

The analysis of the chi-square residuals (sup. Fig S6) shows MITEs and TIRs are the only orders presenting a relative lack of non-polymorphic TEs. Hence, in addition to being the most abundant in the genome, these two TE orders are significantly enriched among polymorphic loci. MITEs are over-represented in both TE polymorphisms types (polymorphic ref. loci and neo-insertions, Fig5 B and D), suggesting a variety of activities within this order. On the other hand, TIRs are found in excess in ref-polymorphisms but lack in neo-insertions. This lack of neo-insertions in TIRs may indicate a recent lower activity in this order, or a more efficient negative selection.

Finally, we observed a strong excess of Maverick among the extra-detection as almost 70% of Maverick polymorphisms (16/23) (Fig 6) fell into this category. Consistent with the observation that, globally, >50% of the extra detections were actually draft annotations eliminated afterwards during filtering steps; ¾ (12/16) of the Maverick elements were also actually present in the draft annotations but later eliminated during filtering steps.

Overall, in proportion, MITEs and TIRs elements are significantly over-represented in TE-polymorphisms. This observation suggests TEs from MITE and TIR orders, in addition to being the most numerous canonical TEs, might have been more active in the genome of *M. incognita* than elements from other TE-orders.

### Some polymorphic loci with contrasted frequency variations between isolates most probably represent true neo-insertions

We investigated the variability of TE presence frequency per locus between the 12 isolates for all the categorized polymorphic loci in the genome.

In ∼ 3/4 (1,911/2,584) of the categorized polymorphic TE loci, the TE presence frequency is homogeneous between isolates (see methods; sup. Fig S8). Said differently, it means that although we observe variations in frequencies between isolates above the estimated error rate (<1%), these variations remain at low amplitude (maximum frequency variation between isolates ≤25% for a given locus). The vast majority (97.95%; 1,872/1,911) concerns loci where the TE is present at a high frequency in all isolates (> 75%). These loci might be considered as fixed in all the isolates. In the remaining 2.04% (39/1,911), the TE frequency is either between 25 and 50% or between 50 and 75% in all isolates. As expected given our methodology, all the high-frequency loci correspond to ref-polymorphisms while all the intermediate frequency loci belong to extra-detections.

In the 673 remaining polymorphic TE loci, TE frequency is heterogeneous, meaning the frequency difference between at least two isolates is > 25% (median difference = 31.35%). Among the most extreme cases of frequency variation per locus, we identified 33 loci in which the TE is found with high frequencies (> 75%) for some isolate(s) while it is absent or rare (frequency <25 %) in the other(s). These loci will be from now on referred to as HCPTEs standing for “Highly Contrasted Polymorphic TE” loci. Because they are highly contrasted, these loci might represent differential fixation/loss across isolates and will be the focus of the following analyses.

HCPTEs encompass 19 MITE elements, 12 TIRs and 2 LINEs (sup. Table S7). We can also notice that some consensuses are more involved in HCPTEs as two TE consensuses alone are responsible for 72.72% (18/33) of these polymorphisms (one MITE consensus involved in 10 HCPTEs, one TIR consensus involved in 8 HCPTEs).

Interestingly, all the HCPTEs loci correspond to neo-insertions regarding the reference genome, meaning that no TE was annotated in the reference genome at this location and the TE presence frequency is < 1% in the Morelos reference isolate. As described in Fig. 7, most of these fixed neo-insertions (20/33) are specific to an isolate and most probably represent lineage-specific neo insertions rather than multiple independent losses.

**Fig 7:**
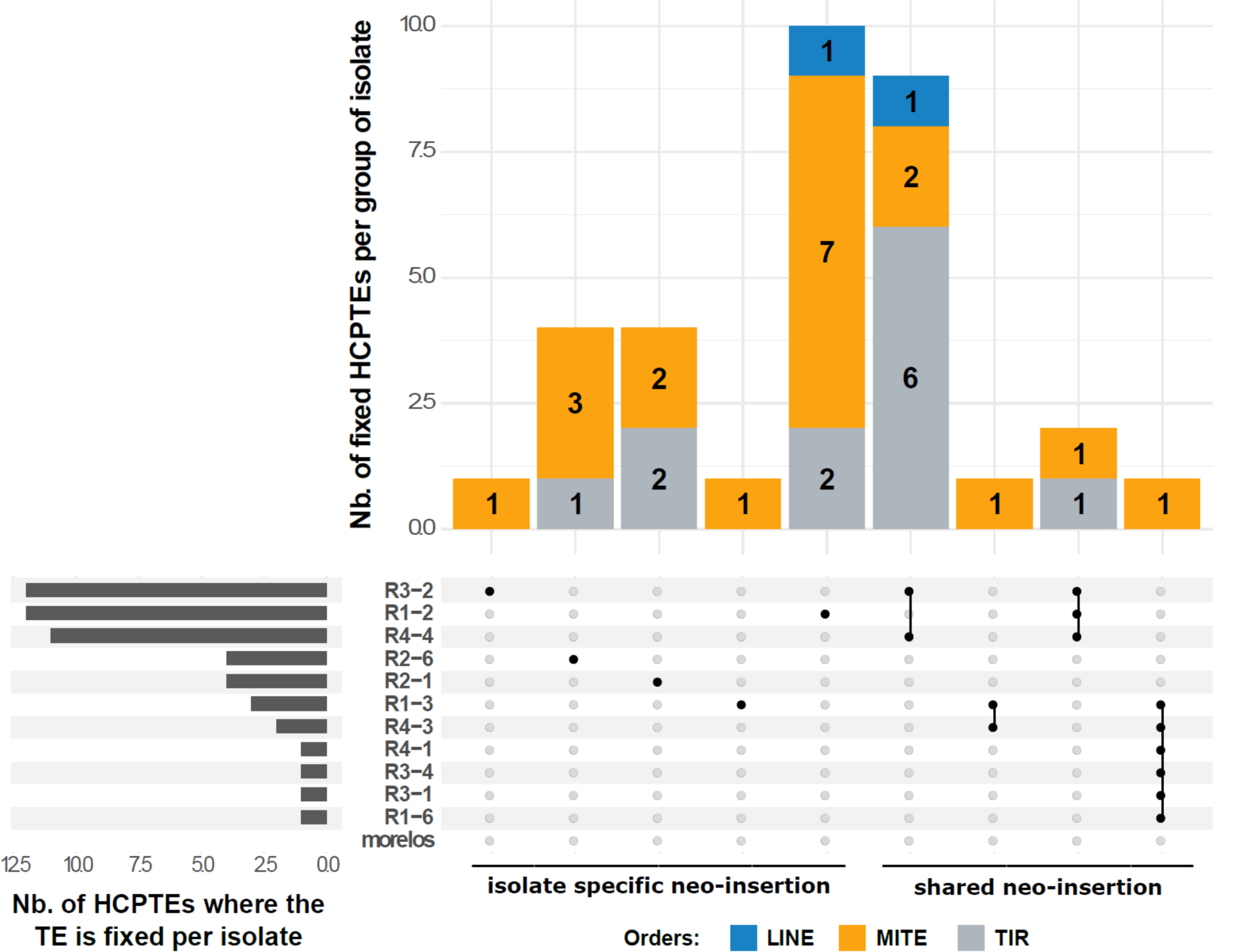
HCPTEs Neo-insertions specificity among the isolates. The central plot shows how many and which isolate(s) share common HCPTEs neo-insertion(s), every line representing an isolate. Columns with several dots linked by a line indicate shared HCPTEs neo-insertion(s) between isolates. Each dot represents which isolate is involved. Columns with a single dot design isolate-specific HCPTEs neo-insertion(s). The top bar plot indicates how many HCPTEs neo-insertions the corresponding group of isolate shares. The left side barplot specifies how many HCPTEs neo-insertion(s) occurred in a given isolate.

However, we also found neo-insertions shared by two (10/33), three (2/33) or even six isolates (1/33). Interestingly, all the shared neo-insertions were between isolates present in a same cluster in the phylogenetic trees (TE-based and SNV-based in Fig. 4), suggesting they might have been fixed in a common ancestor and then inherited. For example, two neo-insertions are shared by isolates R4-4, R1-2 and R3-2 which belong to the same cluster 1 and one neo-insertion is shared by isolates R4-3 and R1-3 which belong to the same cluster 2. Even the neo-insertion shared by 6 isolates follows this pattern as all the concerned isolates belong to the same super-cluster composed of the cluster 2 and 3 plus isolate R1-6 (dashed line in Fig 4).

Hence, the phylogenetic distribution reinforces the idea that these cases are more likely to represent branch-specific neo-insertions than multiple independent losses, including in the reference isolate Morelos.

Isolates R1-2, R3-2, and R4-4 show the highest number of neo-insertions. However, their profiles are quite different. In R1-2, 10/12 HCPTEs are isolate-specific while most of the HCPTEs involving R3-2 and R4-4 are neo-insertions shared with closely related isolates. This is also consistent with the topology and branch lengths of the SNV-based and TE-based phylogenies (sup. Fig S5), which shows that R1-2 is the most divergent isolate with the longest branch length, while R3-2 is quite close to R4-4 and has a relatively short branch.

### Functional impact of TE neo-insertion and validation of in silico predictions

Interestingly, two-thirds (22/33) of the fixed HCPTEs are inserted inside a gene or in a possible regulatory region (1 kb region upstream of a gene). These fixed neo-insertions might have a functional impact in *M. incognita*. Overall, 27 different genes (26 coding for proteins and one tRNA gene) are possibly impacted by the 22 neo-insertions, some genes being in the opposite direction at a neo-insertion point (overlapping this insertion point or being at max 1kb downstream). More than 80% of these genes (22/27) show a substantial expression level during at least one life stage of the nematode life cycle (in the Morelos isolate), suggesting the impacted genes are functional in the *M. incognita* genome (see methods). Some of the impacted genes (40.74%, 11/27) are specific to the *Meloidogyne* genus (they have no predicted orthologs in other nematodes, according to WormBase Parasite). Ten of these Meloidogyne-specific genes are widely conserved in multiple Meloidogyne species, reinforcing their possible importance in the genus, and one is so far only present in *M. incognita*. Interestingly, further similarity search using BLASTp against the NCBI’s nr library returned no significant hits, suggesting these proteins are so far Meloidgyne-specific and do not originate from horizontal gene transfers of non-nematoda origin. Among the remaining genes, one is present in multiple *Meloidogyne* species and otherwise only found in other Plant Parasitic Nematodes species (PPN) (*Ditylenchus destructor, Globodera rostochiensis*) (sup. Table S8). Conservation of these genes across multiple PPN but exclusion from the rest of the nematodes or other species suggest these genes might be involved in important functions relative to these organisms’ lifestyle, including plant parasitism itself.

To experimentally validate *in-silico* predictions of TE neo-insertions with potential functional impact, we performed PCR experiments on 5 of the 22 HCPTEs loci falling in coding or possible regulatory regions (see methods for selection criteria). To perform these PCR validations, we used the DNA remaining from previous extractions performed on the *M. incognita* isolates for population genomics analysis (Koutsovoulos et al. 2020). Basically, the principle was to validate whether the highly contrasted frequencies (>75% / <25%) obtained by PopoolationTE2 actually corresponded to absence/presence of a TE at the locus under consideration (see methods). One isolate (R3-1) presented no amplification in any of the tested loci nor in the positive control. After testing the DNA concentration in the sample, we concluded that the DNA quantity was too low in this isolate and decided to discard it from the analysis.

For four of the five tested HCPTEs loci, we could validate by PCR the *in-silico* predicted differential presence/absence of a sequence at this position, across the different isolates (Fig 8; (Kozlowski, Hassanaly-Goulamhoussen, et al. 2020)).

**Fig 8:**
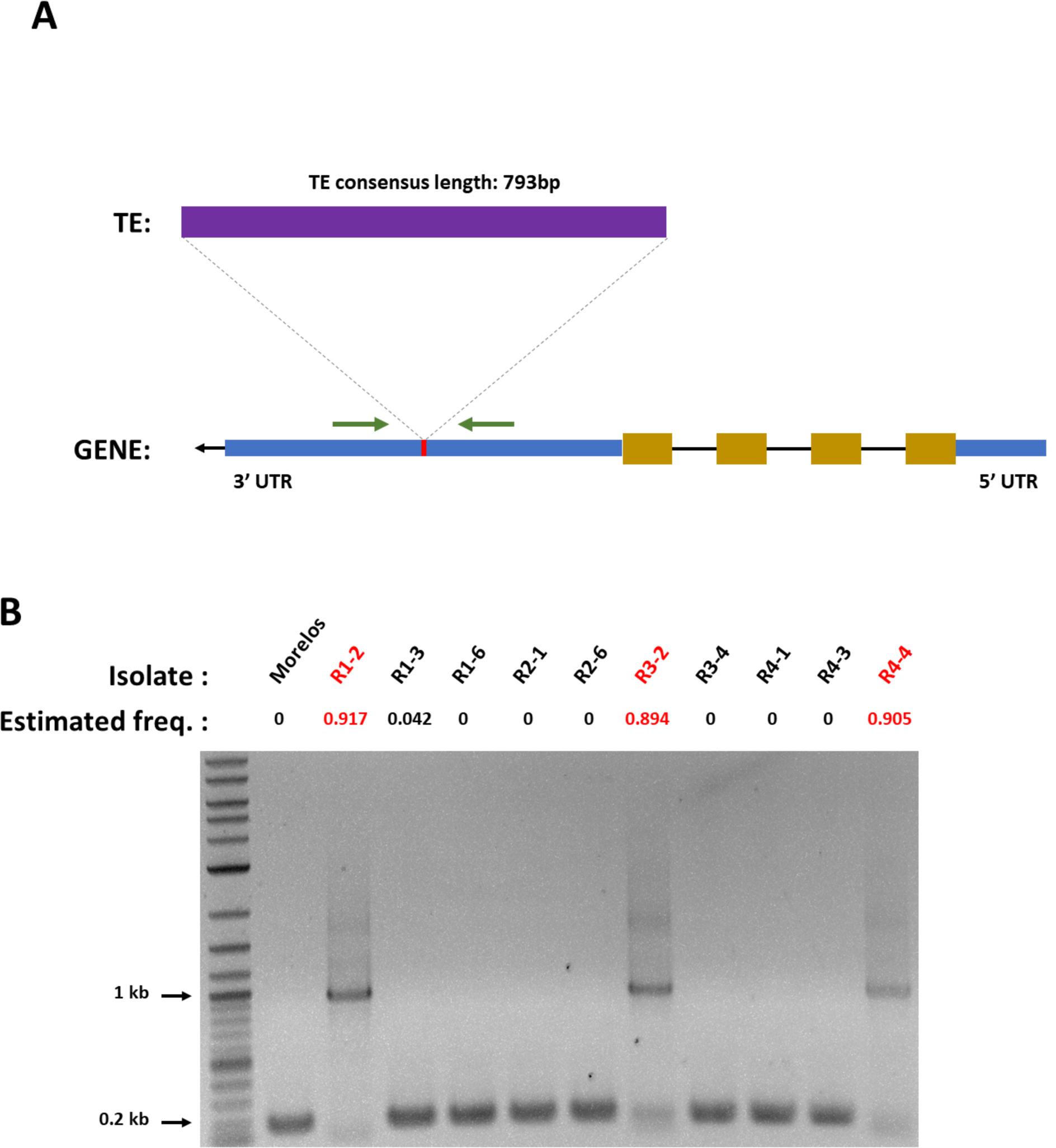
Experimental validation of a predicted neo-insertion. A-Diagram of the TE neo-insertion. The neo-insertion of the MITE element occurs in the 3’UTR region of the gene (Minc3s00026g01668). Blue boxes illustrate the 3’ and 5’ UTR regions of the gene while the yellow boxes picture the exons. Green arrows represent the primers used to amplify the region. Gene subparts and TE representations are not at scale. Predicted size of the amplicon: 973 bp with the TE insertion, 180 bp without. B-PCR validation of the TE neo-insertion. Estimated freq. values correspond to the proportion of individuals per isolate predicted to have the TE at this position (PopoolationTE2). Isolates in red were predicted to have the TE inserted at this locus. Only these isolates show an amplicon with a size suggesting an insertion (sequences are available in (Kozlowski, Hassanaly-Goulamhoussen, et al. 2020)).

In one of the five tested loci, named locus 1, we could i) validate by PCR the presence of a sequence at this position for the isolates presenting a PopoolationTE2 frequency >75% and absence for those having a frequency <25%; ii) also validate by sequencing that the sequence itself corresponded to the TE under consideration (a MITE). This case is further explained in detail below and in Fig. 8.

According to PopoolationTE2 frequencies, in the concerned locus, 1 MITE is inserted and fixed in 3 isolates (R1-2, R3-2, R4-4) as the estimated frequencies are higher than 75% in these isolates. We assumed the TE is absent from the rest of the isolates as all of them display frequencies <5%. To validate this differential presence across the isolates, we designed specific primers from each side of the estimated insertion point so that the amplicon should measure 973 bp with the TE insertion and 180 bp without.

The PCR results are consistent with the frequency predictions as only R1-2, R3-2, and R4-4 display a ∼1 kb amplicon while all the other isolates show a ∼0.2 kb amplicon (Fig 8). Hence, as expected, only the 3 isolates with a predicted TE frequency >75% at this locus exhibit a longer region, compatible with the MITE insertion.

To validate the amplified regions corresponded to the expected MITE, we sequenced the amplicons for the 3 predicted insertions and aligned the sequences to the TE consensus and the genomic region surrounding the estimated insertion point (Kozlowski, Hassanaly-Goulamhoussen, et al. 2020). Amplicon sequences of R-1_2, R-3_2, and R-4_4 all covered a significant part of the TE consensus sequence length (> 78%) with high % identity (> 87%) and only a few gaps (<5%). These results confirm that the inserted sequence corresponds to the predicted TE consensus. Moreover, all the 3 amplicons aligned on the genomic region downstream of the insertion point with high % identity (>= 99%), which helped us further determine the real position of the insertion point. The real insertion point is 26 bp upstream of the one predicted by PopoolationTE2 and falls in the forward primer sequence. This explains why the amplicon sequences do not align on the region upstream the insertion point.

We also noticed that the inserted TE sequences slightly diverged between the isolates while the genomic region surrounding the insertion point remains identical. Interestingly, the level of divergence in the TE sequence does not follow the phylogeny as R-4_4 is closer to R-1_2 than to R-3_2 (sup. Table S9).

Finally, in the Morelos, R-2_1, and R-2_6 isolates, the sequencing of the amplicon validated the absence of insertions. Indeed, the sequences aligned on the genomic region surrounding the insertion point with high % identity (99, 97, 87 % respectively) but not with the MITE consensus.

Hence, we fully validated experimentally the presence/absence profile across isolates predicted *in silico* at this locus.

In the *M. incognita* genome, this neo-insertion is predicted to occur in the 3’ UTR region of a gene (Minc3s00026g01668). This gene has no obvious predicted function, as no conserved protein domain is detected and no homology to another protein with an annotated function could be found. However, orthologs were found in the genomes of several other *Meloidogyne* species (*M. arenaria, M. javanica, M. floridensis, M. enterolobii*, and *M. graminicola*), ruling out the possibility that this gene results from a prediction error from gene calling software. The broad conservation of this gene in the *Meloidogyne* genus suggests this gene might be important for *Meloidogyne* biology and survival.

In the Morelos isolate, for which no TE was inserted at this position, this gene is supported by transcriptomic RNA-seq data during the whole life cycle of the nematode (Kozlowski, Da Rocha, et al. 2020), suggesting this gene is probably functionally important in *M. incognita* and other root-knot nematodes.

Consequently, the insertion of the TE in R-1_2, R-3_2, and R-4_4 genome at this locus could have functional impacts.

## Discussion

### TE landscape in nematode genomes and possible recent activity in *M. incognita*

In this analysis, we have annotated TEs in the genome of *M. incognita* and used variations in TE frequencies between geographical isolates across loci as a reporter of their activity. The *M. incognita* TE landscape is more abundant in DNA than retro-transposons and using the same methodology, we confirmed a similar trend in the genome of *C. elegans*. Interestingly, even if the methodology used was different, a similar observation was made at the whole nematoda level (Szitenberg et al., 2016), suggesting a higher abundance of DNA transposons might be a general feature of nematode genomes.

We have shown 75% of the polymorphic TE loci in *M. incognita* display moderate frequency variations between isolates (<25%); a majority being found with high frequencies (> 75%) in all the isolates simultaneously. Hence, a substantial part of the TE can be considered as stable and fixed among the isolates.

Nevertheless, the remaining quarter of polymorphic TE loci present frequency variations across the isolates exceeding 25%. This observation concerns both the TE already present in the reference genome, but also the neo-insertions. We even detected loci where the TE frequencies were so contrasted between the isolates (HCPTEs) that we could predict the TE presence/absence pattern among the isolates. Such frequency variations between isolates, and the fact that part of the HCPTEs are isolate-specific neo-insertions, constitute strong evidence for TE activity in the *M. incognita* genome.

In *C. elegans*, multiple TE families have also shown a substantial level of activity across different populations (Laricchia et al. 2017). However, this analysis was based on binary presence / absence data of TE at loci across populations and thus provided no information about the amplitude of TE frequencies variability within isolates. In our analysis we provided this extra layer of information and this also allowed estimating the amplitude of TE frequency variations between *M. incognita* isolates.

It should be noted here that the total TE activity in the *M. incognita* genome is probably underestimated, in part because of our strategy to eliminate false positives as much as possible by applying a series of stringent filters, and in another part because of the intrinsic limitations of the tools, such as the incapacity of PopoolationTE2 to detect nested TEs (Kofler et al. 2016).

We then evaluated how recent this activity could be, using % identity of the TE copies with their respective consensuses as a proxy for their age as previously proposed in other studies (Bast et al. 2015; Lerat et al. 2019). We showed that a substantial proportion of the canonical TE annotations were highly similar to their consensus, indicating most of these TE copies were recent in the genome. The probable recent hybrid origin of *M. incognita* (Blanc-Mathieu et al. 2017) is consistent with a recent TE burst in the genome. Indeed, as further explained in the last section of the discussion, it is well established that hybridization events can lead to a relaxation of the TE silencing mechanisms and consequently to a TE expansion (Belyayev 2014; Guerreiro 2014; Rodriguez and Arkhipova 2018).

However, as suggested in (Bourgeois and Boissinot 2019), the extent of this phenomenon might differ depending on the TE order. In *M. incognita*, MITEs and TIRs alone account for ∼2/3 of the canonical TE annotations, but their fate in the genome seems to have followed different paths. Indeed, as illustrated in Figure 2, MITEs show a wide range of identity rate with their consensus, which suggests they might have progressively invaded the genome being uncontrolled or poorly controlled as suggested for the rice genome (Lu et al. 2017). On the opposite, almost all the TIR copies share high percentage identity with their consensuses which could be reminiscent of a rapid and recent burst. Nevertheless, this burst could have quickly been under control as, according to chi-square residuals (sup. Fig S6), TIR neo-insertions are significantly less numerous than expected owing to their abundance in the genome. Interestingly, in *C. elegans*, the Tc1 / Mariner TIR DNA element was shown to be the most active while, so far, no evidence for active retro-transposition was shown in this species (Bessereau 2006; Laricchia et al. 2017).

Because no molecular clock is available for *M. incognita*, it is impossible to evaluate more precisely when TE bursts would have happened and how fast each TE from each order would have spread in the genome. Such bursts can be very recent, including in animal genomes as exemplified by the P-element which invaded the genome of some Drosophila populations in just 40 years (Anxolabéhère et al. 1988). While an absolute dating of TE activities in *M. incognita* is currently not possible, a relative timing of the events regarding population diversification can still be deduced from the distribution of TE loci frequencies across isolates. Indeed, we have shown (Figure 7) that some neo-insertion were shared between isolates and that in each case, the concerned isolates belonged to a same monophyletic cluster (Figure 4). The most parsimonious scenario is that these neo-insertions occurred in *M. incognita*, after the separation of the different main clusters but before the diversification of the phylogenetically-related isolates, within a cluster, in a common ancestor. Other TE neo-insertions, in contrast, were so far isolate-specific, suggesting some TE movements were even more recent and that TE mobility might be a continuous phenomenon. No information is available about the ancientness of cultivated lands in Brazil on which the different isolates have been sampled. However, because there is no significant correlation between the isolates geographical distribution and the phylogenetic clusters, whether it is TE-based (this study) or SNV-based (Koutsovoulos et al. 2020), we can hypothesize these isolates have been recently spread by human agricultural activity in the last centuries.

Overall, the presence of isolate-specific TE neo-insertions, the distribution of percent identities of some TE copies to their consensuses shifted towards high value, as well as transcriptional support for some of the genes involved in the transposition machinery, suggest TE have recently been active in *M. incognita* and are possibly still active.

### Functional impact of TEs activity in *M. incognita* and other nematodes

*M. incognita* is a parthenogenetic mitotic nematode of major agronomic importance. How this pest adapts to its environment in the absence of sexual recombination remains unresolved. In this study, we investigated whether TE movements could constitute a mechanism of genome plasticity compatible with adaptive evolution.

In *M. javanica*, a closely related root-knot nematode, comparison between an avirulent line unable to infect tomato plants carrying a nematode resistance gene and another virulent line that overcame this resistance, led to the identification of a gene present in the avirulent nematodes but absent from the virulent ones. Interestingly, the gene under consideration is present in a TIR-like DNA transposon and its absence in the virulent line suggests this is due to excision of the transposon and thus that TE activity plays a role in *M. javanica* adaptive evolution (Gross and Williamson 2011).

In *M. incognita*, convergent gene losses at the whole genome level between two virulent populations compared to their avirulent populations of origin were recently reported (Castagnone-Sereno et al. 2019). Gene copy number variation CNV are indeed known to be involved in genomic plasticity and in adaptive evolution (Katju and Bergthorsson 2013), and TE can actively (e.g. by gene hitchhiking) or passively (e.g. through illegitimate recombination) participate in these variations. This CNV analysis in *M. incognita* was done on an older version of the genome (Abad et al. 2008), that was partially incomplete, and the possible contribution of TEs in these CNV could not be assessed. Although the current version of the genome (Blanc-Mathieu et al. 2017) is more complete and consistent with the estimated genome size, it is still fragmentary with thousands of scaffolds and a relatively low N50 length (38.6 kb). This fragmentation prevents a thorough identification of TE-rich and TE-poor regions and possible co-localization with CNV loci at the whole genome scale. Availability of long read-based more contiguous genome assembly in the future will certainly allow reinvestigating CNV and the possible involvement of TEs in association to an adaptive process such as resistance breaking down.

As previously evoked, in *M. incognita*, we found that the genome-wide pattern of variations of TE frequencies across the loci between the different populations recapitulated almost exactly the phylogeny of the isolates built on SNV in coding regions (Fig 4). Hence, most of the divergence in terms of TE pattern follows the divergence at the nucleotide level and thus the phylogeny of the isolates. Almost the same conclusion was drawn by comparing SNV and TE variation data across different *C. elegans* populations (Laricchia et al. 2017). In *M. incognita*, the phylogeny of isolates does not significantly correlate with the monitored biological traits, namely geographical distribution, range of compatible host plants and nature of the crop currently infected (Koutsovoulos et al. 2020). Interestingly, no correlation was also observed between variations in TE frequencies and geographical distribution for European *Drosophila* populations (Lerat et al. 2019). The lack of evident correlation between the phylogenetic signal regardless whether it is TE-based or SNV-based and the biological traits under consideration suggests most of the variations follow the drift between isolates and are not necessarily adaptive, which is not surprising. A similar conclusion was also drawn recently by analyzing 625 fungal genomes and observing that most TE movements were presumably neutral and adaptive ones being marginal (Muszewska et al. 2019).

On another note, as explained in the first section of the discussion, TE activity is possibly very recent in *M. incognita* and this might contribute to the current lack of evidence for association between TE activity, including invasion or decay across populations and adaptive traits.

Yet, we detected, and confirmed by PCR the neo-insertions of TE in some functionally important loci, inside genes or possible regulatory regions. We found that more than 90% of the TEs involved were TIRs or MITEs, which echoes their enrichment among the most active TEs in *M. incognita*. In the Mulberry genome, MITEs inserted near genes were shown to regulate gene expression via small RNAs while those inserted within genes were associated with alternative splice variants (Xin et al. 2019). Similarly, in the wheat genome, MITEs of the mariner superfamily played an instrumental role in generating the diversity of micro-RNAs involved in important adaptive traits such as resistance to pathogens (Poretti et al. 2020). The exact functional impact of TE insertions in *M. incognita* would need to be evaluated in the future. Generating transcriptomics data for the different isolates would enable studying associated differences in gene expression patterns or transcript diversity. As a complementary approach, proteomic studies would allow direct search for differences at the encoded protein level.

Regardless of the future experimental validation of the functional impact, one important question concerns the current preliminary evidence for a possible role in the nematode adaptive evolution. Because some of the impacted genes are specific to plant-parasitic species and yet conserved in several of these phyto-parasites, a role in plant parasitism is possible. Interestingly, TE movements can be involved in the emergence of species or genus-specific ‘orphan’ genes (Ruiz-Orera et al. 2015; Wu and Knudson 2018; Jin et al. 2019). However, in the absence of known protein domains or functional characterization of these genes, the exact biochemical activity or biological processes in which they might be involved remains elusive.

### Ploidy, (a)sexuality and hybridization: a complex interplay on TE load and composition

*M. incognita* is an asexual (mitotic parthenogenetic), polyploid, and hybrid species. These three features are expected to impact TE load in the genome with various intensities and possibly conflicting effects.

Contradictory theories exist concerning the activity/proliferation of TEs as a function of the reproductive mode. The higher efficacy of selection under sexual reproduction can be viewed as an efficient system to purge TEs and control their proliferation. Supporting these views, in parasitoid wasps, TE load was shown to be higher in asexual lineages induced by the endosymbiotic Wolbachia bacteria than in sexual lineages (Kraaijeveld et al. 2012). However, whether this higher load is a consequence of the shift in reproductive mode or of Wolbachia infection remains to be clarified.

In an opposite theory, sexual reproduction can also be considered as a way for TEs to spread across individuals within the population whereas in clonal reproduction the transposons are trapped exclusively in the offspring of the holding individual. Under this view, asexual reproduction is predicted to reduce TE load as TE are unable to spread in other individuals, and are thus removed by genetic drift and/or purifying selection in the long term (Wright and Finnegan 2001). Consistent with this theory, comparison of sexual and asexual *Saccharomyces cerevisiae* populations showed that the TE load decreases rapidly under asexual reproduction (Bast et al. 2019).

Hence, whether the TE-load is expected to be higher or lower in clonal species compared to sexual relatives remains unclear and other conflicting factors such as TE excision rate and the effective size of the population probably blur the signal (Glémin et al. 2019). The breeding system has been shown to constitute an important factor,of TE distribution in Caenorhabditis genomes (Dolgin et al. 2008): TEs in self-fertilizing populations seem to be selectively neutral and segregate at higher frequency than in outcrossing populations, where they are submitted to purifying selection. Interestingly, at a broader scale, a comparative analysis of different lineages of sexual and asexual arthropods revealed no evidence for differences in TE load according to the reproductive modes (Bast et al. 2015). Similar conclusions were drawn at the whole nematoda phylum scale (Szitenberg et al. 2016), although only one apomictic asexually-reproducing species (i.e. *M. incognita*) was present in the comparative analysis.

Polyploidy, in contrast, is commonly accepted as a major event initially favouring the multiplication and activity of TEs. This is clearly described with numerous examples in plants (Vicient and Casacuberta 2017) and some examples are also emerging in animals (Rodriguez and Arkhipova 2018). When hybridization and polyploidy are combined, this can lead to TE bursts in the genome. As originally proposed by Barbara McClintock, allopolyploidization produces a “genomic shock”, a genome instability associated with the relaxation of the TE silencing mechanisms and the reactivation of ancient TEs (McClintock 1984; Mhiri et al. 2019).

Hybridization, polyploidy and asexual reproduction are combined in *M. incognita* with relative effects on the TE load extremely challenging, if not impossible, to disentangle. Initial comparisons of the TE loads in three allopolyploid clonal *Meloidogyne* against a diploid facultative sexual relative suggested a higher TE load in the clonal species (Blanc-Mathieu et al. 2017). However, to differentiate the relative contribution of each of these three features to the *M. incognita* TE load, it would be necessary to conduct comparative analysis with a same method on diploid asexuals, on polyploid sexuals as well as on diploid asexuals in the genus *Meloidogyne*, and ideally with and without hybrid origin. So far, genomic sequences are only available for other polyploid clonal species, which are all suspected to have a hybrid origin (Blanc-Mathieu et al. 2017; Szitenberg et al. 2017; Koutsovoulos et al. 2019; Susič et al. 2020), and, apart from that, only two diploid facultative sexual species (Opperman et al. 2008; Somvanshi et al. 2018). Hence, further sampling of *Meloidogyne* species with diverse ploidy levels and reproductive modes will be necessary to disentangle the relative contribution of ploidy level, hybridization and reproductive mode on the TE abundance and composition.

## Concluding remarks

In this study we used population genomics technique and statistical analyses of the results to assess whether TE might contribute to the genome dynamics of *M. incognita* and possibly to its adaptive evolution. Overall, we provided a body of evidence suggesting TE have been at least recently active and might still be active. With thousands of loci showing variations in TE presence frequencies across geographical isolates, there is a clear impact on the *M. incognita* genome plasticity. Some TE being neo-inserted in coding or regulatory regions might have a functional impact. Although no clear connection with a role in adaptive evolution could be made so far, based on the few impacted coding loci we experimentally checked in this study, this is not to be excluded given the current lack of large-scale functional information for this species. This pioneering study constitutes a valuable resource and opens new perspectives for future targeted investigation of the potential effect of TE dynamics on the evolution, fitness and adaptability of *M. incognita* as well as in the whole nematoda phylum.

## Materials and Methods

### Material

#### The genome of *M. incognita*

We used the genome assembly published in (Blanc-Mathieu et al. 2017) as a reference for TE prediction and annotation (ENA assembly accession GCA_900182535, bioproject PRJEB8714) as well as for read-mapping of the different geographical isolates (Koutsovoulos et al. 2020), used for prediction of TE presence frequencies.

Briefly, the triploid *M. incognita* genome is 185Mb long with ∼12,000 scaffolds and a N50 length of ∼38 kb. Although the genome is triploid, because of the high nucleotide divergence between the genome copies (8% on average), most of these genome copies have been correctly separated during genome assembly, which can be considered effectively haploid (Blanc-Mathieu et al. 2017; Koutsovoulos et al. 2020). This reference genome originally came from a *M. incognita* population from the Morelos region of Mexico and was reared on tomato plants from the offspring of one single female in our laboratory.

#### The genome of *C. elegans*

We used the *C. elegans* genome (The C. elegans Genome Sequencing Consortium 1998) assembly (PRJNA13758) to perform its repeatome prediction and annotation and compare our results to the literature as a methodological validation.

#### Genome reads for 12 *M. incognita* geographical isolates

To predict the presence frequencies at TE loci across different *M. incognita* isolates, we used whole-genome sequencing data from pools of individuals from 12 different geographical regions (sup. Fig S4 & sup. Table S10). One pool corresponds to the Morelos isolates used to produce the *M. incognita* reference genome itself, as described above. The 11 other pools correspond to different geographical isolates across Brazil as described in (Koutsovoulos et al. 2020).

All the samples were reared from the offspring of one single female and multiplied on tomato plants. Then, approximately 1 million individuals were pooled and sequenced by Illumina paired-end reads (2*150bp). Libraries sizes vary between 74 and 76 million reads (Koutsovoulos et al. 2020).

We used cutadapt-1.15 (Martin 2011) to trim adapters, discard small reads, and trim low-quality bases in reads boundaries (–max-n=5 -q 20,20 -m 51 -j 32 -a AGATCGGAAGAGCACACGTCTGAACTCCAGTCA -A AGATCGGAAGAGCGTCGTGTAGGGAAAGAGTGT). Then, for each library, we performed a fastqc v-0.11.8 (Andrew S., 2010: http://www.bioinformatics.babraham.ac.uk/projects/fastqc) analysis to evaluate the quality of the reads. FastQC results analyses showed that no additional filtering or cleaning step was needed and no further read was discarded.

### Methods

We performed the statistical analysis and the graphical representation using R’ v-3.6.3 and the following libraries: ggplot2, cowplot, reshape2, ggpubr, phangorn, tidyverse, and ComplexUpset. All codes and analysis workflows are publicly available in the INRAE Dataverse (Kozlowski 2020a; Kozlowski 2020c; Kozlowski, Da Rocha, et al. 2020). For experimental validations, see (Kozlowski, Hassanaly-Goulamhoussen, et al. 2020). A diagram recapitulating the main steps of the analysis has been provided in supplementary (sup. Fig S7); as well as a decision tree summarising the polymorphism characterisation (sup. Fig S8).

#### *M. incognita* and *C. elegans* repeatome predictions and annotations

We predicted and annotated the *M. incognita* and *C. elegans* repeatomes following the same protocol as thoroughly explained in (Koutsovoulos et al. 2019). We define the repeatome as all the repeated sequences in the genome, excluding Simple Sequence Repeats (SSR) and microsatellites. Then, following the above-mentioned protocol, we further analysed each repeatome to isolate annotations with canonical signatures of Transposable Elements (TEs).

Below, we briefly explain each step and describe protocol adjustments.

##### Genome pre-processing

Unknown nucleotides ‘Ns’ encompass 1.81% of *the M. incognita* reference genome and need to be trimmed before repeatome predictions. We created a modified version of the genome by splitting it at N stretches of length 11 or more and then trimming all N, using dbchunk.py from the REPET package (Quesneville et al. 2005; Flutre et al. 2011). As this increases genome fragmentation and may, in turn, lead to false positives in TE detection, we only kept chunks of length above the L90 chunk length threshold, which is 4,891 bp. This modified version of the genome was only used to perform the *de novo* prediction of the TE consensus library (below). The TE annotation was performed on the whole reference genome.

The *C. elegans* reference genome was entirely resolved (no N), at the chromosome-scale. Hence, we used the whole assembly as is to perform the *de novo* prediction analysis.

##### De novo prediction: constituting draft TE-consensus libraries

For each species, we used the TEdenovo pipeline from the REPET package to generate a draft TE-consensus library..

Briefly, TEdenovo pipeline i) realises a self-alignment of the input genome to detect repetitions, ii) clusters the repetitions, iii) performs multiple alignments from the clustered repetitions to create consensus sequences, and eventually, iv) classify the consensus sequence following the Wicker’s classification (Wicker et al. 2007) using structural and homology based information. One of the most critical steps of this process concerns the clustering of the repetitions as it requires prior knowledge about assembly ploidy and phasing quality.

We ran the analysis considering the modified *M. incognita* reference assembly previously described as triploid and set the ‘minNbSeqPerGroup’ parameter to 7 (*i*.*e* 2n+1). As the *C. elegans* assembly was haploid, we set the same parameter to 3.

All the remaining parameters values set in these analyses can be found in the TEdenovo configuration files (Kozlowski 2020a).

##### Automated curation of the TE-consensus libraries

To limit the redundancy in the previously created TE consensus libraries and the false positives, we performed an automated curation step. Briefly, for each species, i) we performed a minimal annotation (steps 1, 2, 3, 7 of TEannot) of their genome with their respective draft TE-consensus libraries, and ii) only retained consensus sequences with at least one Full-Length Copy (FLC) annotated in the genome. All parameters values are described in the configuration files available in (Kozlowski 2020a).

##### Repeatome annotation

For each species, we performed a full annotation (steps 1, 2, 3, 4, 5, 7, and 8) of their genome with their respective cleaned TE-consensus libraries using TEannot from the REPET package. The obtained repeatome annotations (excluding SSR and microsatellites) were exported for further analyses. All parameters values are described in the configuration files available in (Kozlowski 2020a).

##### Repeatome post-processing: identifying annotations with canonical signatures of TEs

Using in house scripts (Kozlowski 2020a), we analysed REPET outputs to retain annotations with canonical signatures of Transposable Elements (TEs) from the rest of the repeatomes. The same parameters were set for *M. incognita* and *C. elegans*. Briefly, for each species, we only conserved TE annotations i) classified as retro-transposons or DNA-transposons, ii) longer than 250 bp, iii) sharing more than 85% identity with their consensus sequence, iv) covering more than 33% of their consensus sequence length, v) first aligning with their consensus sequence in a BLAST analysis against the TE-consensus library, and vi) not overlapping with other annotations. TE annotations respecting all the described criterion were referred to as canonical TE annotations.

#### Putative transposition machinery identification (*M. incognita* only)

We analysed the *M. incognita* predicted proteome and transcriptome (Blanc-Mathieu et al. 2017) and crossed the obtained information with the canonical TE-annotation to identify TE containing genes putatively involved in the transposition machinery and evaluate TE-related gene expression levels in comparison to the rest of the genes in the genome.

##### Finding genes coding for proteins with TE-related HMM profiles

We performed an exhaustive HMMprofile search analysis on the whole *M. incognita* predicted proteome and then looked for proteins with TE-related domains. First, we concatenated two HMMprofile libraries into one: Pfram32 (Finn et al. 2016) library and Gypsy DB 2.0 (Llorens et al. 2011), a curated library of HMMprofiles linked to viruses, mobile genetic elements, and genomic repeats. Then, using this concatenated HMM profile library, we performed an exhaustive but stringent HMM profile search on the *M. incognita* proteome using hmmscan (-E 0.00001 --domE 0.001 --noali).

Eventually, using in house script (Kozlowski, Da Rocha, et al. 2020), we selected the best non-overlapping HMM profiles for each protein and then tagged corresponding genes with TE-related HMM profiles thanks to a knowledge-based function from the REPET tool ‘profileDB4Repet.py’. We kept as genes with TE-related profiles all the genes with at least one TE-related HMM-profile identified.

##### Genes expression level

To determine the *M. incognita* protein-coding genes expression patterns, we used data from a previously published life-stage specific RNA-seq analysis of *M. incognita* transcriptome during tomato plant infection (Blanc-Mathieu et al. 2017). This analysis encompassed four different life stages: (i) eggs, (ii) pre-parasitic second stage juveniles (J2), (iii) a mix of late parasitic J2, third stage (J3) and fourth stage (J4) juveniles and (iv) adult females, all sequenced in triplicates.

The cleaned RNA-seq reads were retrieved from the previous analysis and re-mapped to the *M. incognita* annotated genome assembly (Blanc-Mathieu et al. 2017) using a more recent version of STAR (2.6.1) (Dobin et al. 2013) and the more stringent end-to-end option (*i*.*e*. no soft clipping) in 2-passes. Expected read counts were calculated on the predicted genes from the *M. incognita* GFF annotation as FPKM values using RSEM (Li and Dewey 2011) to take into account the multi-mapped reads via expectation maximization. To reduce amplitude of variations, raw FPKM values were transformed to Log10(FPKM+1) and the median value over the 3 replicates was kept as a representative value in each life stage. The expression data are available in (Danchin and Da Rocha 2020).

Then, for each life stage independently, i) we ranked the gene expression values, and ii) defined gene expression level corresponding to the gene position in the ranking. We considered as substantially expressed all the genes that presented an expression level >= 1st quartile in at least one life stage.

##### TE annotations with potential transposition machinery

To identify TE-annotations including predicted genes involved in transposition machinery (inclusion >= 95% of the gene length), we performed the intersection of the canonical TE annotation and the genes annotation BED files (Kozlowski, Da Rocha, et al. 2020) using the intersect tool (-wo -s -F 0.95) from the bedtools v-2.27.1 suite (Quinlan and Hall 2010).

We then cross-referenced the obtained file with the list of the substantially expressed genes and the list of the TE-related genes previously elaborated to identify the TEs containing potential transposition machinery genes and their expression levels.

#### Evaluation of TE presence frequencies across the different *M. incognita* isolates

We used the popoolationTE2 v-1.10.04 pipeline (Kofler et al. 2016) to compute isolate-related support frequencies of both annotated, and *de novo* TE-loci across the 12 *M. incognita* geographical isolates previously described. To that end, we performed a ‘joint’ analysis as recommended by the popoolationTE2 manual. Briefly, popoolationTE2 uses both quantitative and qualitative information extracted from paired-end (PE) reads mapping on the TE-annotated reference genome and a set of reference TE sequences to detect signatures of TE polymorphisms and estimate their frequencies in every analysed isolate. Frequency values correspond to the proportion of individuals in an isolate for which a copy of the TE is present at a given locus.

##### Preparatory work: creating the TE-hierarchy and the TE-merged-reference files

We used the canonical TE-annotation set created above (Kozlowski 2020a) and the *M. incognita* reference genome to produce the TE-merged reference file and the TE-hierarchy file necessary to perform the popoolationTE analysis (Kozlowski 2020c).

We used getfasta and maskfasta commands (default parameters) from the bedtools suite to respectively extract and mask the sequences corresponding to canonical TE-annotations in the reference genome. Then we concatenated both resulting sequences in a ‘TE-merged reference’ multi fasta file. The ‘TE-hierarchy’ file was created from the TE-annotation file from which it retrieves and stores the TE sequence name, the family, and the TE-order for every entry.

##### Reads mapping

For each *M. incognita* isolate library, we mapped forward and reverse reads separately on the “TE-merged-references” genome-TE file using the local alignment algorithm bwa bwasw v-0.7.17-r1188 (Li and Durbin 2009) with the default parameters. The obtained sam alignment files were then converted to bam files using samtools view v-1.2 (Li et al. 2009).

##### Restoring paired-end information and generating the ppileup file

We restored paired-end information from the previous separate mapping using the sep2pe (--sort) tool from popoolationTE2-v1.10.03. Then, we created the ppileup file using the ‘ppileup’ tool from popolationTE2 with a map quality threshold of 15 (--map-qual 15).

For every base of the genome, this file summarises the number of PE reads inserts spanning the position (physical coverage) but also the structural status inferred from paired-end read covering this site.

##### Estimating target coverage and subsampling the ppileup to a uniform coverage

As noticed by R. Kofler, heterogeneity in physical coverage between populations may lead to discrepancies in TE frequency estimation. Hence, we flattened the physical coverage across the *M. incognita* isolates by a subsampling and a rescaling approach.

We first estimated the optimal target coverage to balance information loss and homogeneity using the ‘stats-coverage’ tool from PopoolationTE2 (default parameter) and set this value to 15X. We then used the ‘subsamplePpileup’ tool (--target-coverage 15) to discard positions with a physical coverage below 15X and rescale the coverage of the remaining position to that value.

##### Identify signatures of TE polymorphisms

We identified signatures of TE polymorphisms from the previously subsampled file using the ‘identifySignature’ tool following the joint algorithm (--mode joint; --min-count 2; --signature-window minimumSampleMedian; --min-valley minimumSampleMedian).

Then, for each identified site, we estimated TE frequencies in each isolate using the ‘frequency’ tool (default parameters). Eventually, we paired up the signatures of TE polymorphisms using ‘pairupSignatures’ tool (--min-distance -200; --max-distance -- 300 as recommended by R. Kofler), yielding a final list of potential TE-polymorphisms positions in the reference genome with their associated frequencies for each one of the isolates.

##### Evaluation of PopoolationTE2 systematic error rate in the TE-frequency estimation

To estimate PopoolationTE2 systematic error rate in the TE-frequency estimation, we ran the same analysis (from the PE information restoration step) but comparing each isolate against itself (12 distinct analyses).

We then analysed each output individually, measuring the frequency difference between the two ‘replicates’ in all the detected loci with FR signatures (see below for more explanations).

We tested the homogeneity of the frequency-difference across the 12 analyses with an ANOVA and concluded that the mean values of the frequencies differences between the analysis were not significantly heterogeneous (p. value = 0.102 > 0.05). Hence, we concatenated the 12 analysis frequency-difference and set the systematic error rate in the TE-frequency estimation to 2 times the standard deviation of the frequency differences, a value of 0.97 %.

#### TE polymorphism analysis

##### Isolating TE loci with frequency variation across M. incognita isolates

We parsed PopoolationTE2 analysis output to identify TE loci with enough evidence to characterise them as polymorphic in frequency across the isolates.

PopoolationTE2 output informs for each detected locus i) its position on the reference genome, ii) its frequency value for every sample of the analysis (e.g each isolate), and iii) qualitative information about the reads mapping signatures supporting a TE insertion.

In opposition to separate Forward (‘F’) or Reverse (‘R’) signatures, ‘FR’ signatures mean the locus both boundaries are supported by significant physical coverage. Entries with such type of signature are more accurate in terms of frequency and position estimation. Hence, we only retained candidate loci with ‘FR’ signatures. Then, for each locus, we computed the maximal frequency variation between all the isolates and discarded the loci with a frequency difference smaller than the PopoolationTE2 systematic error rate in the TE-frequency estimation we computed (0.97 %; see above). We also discarded loci where different TEs were predicted to be inserted. We considered the remaining loci as polymorphic in frequency across the isolates.

##### Isolates phylogeny

We reconstructed *M. incognita* isolates phylogeny according to their patterns of polymorphism in TE frequencies.

We first computed a euclidean distance matrix from the isolates TE frequencies of all the detected polymorphic loci. We then used the distance matrix to construct the phylogenetic tree using the Neighbor Joining (NJ) method (R’ phangorn package v-2.5.5). We computed nodes support values with a bootstrap approach (n=500 replicates) using the boot.phylo function from the ape-v5.4 R package (Paradis and Schliep 2019). The boot.phylo function performs a resampling of the frequency matrix (here the matrix with loci in columns, isolates in row, and values corresponding to the TE presence frequencies).

Also, we created a phylogenetic tree using the SNV from coding regions for all isolates with raxml-ng v-0.9.0 (Kozlov et al. 2019) utilising the model GTR+G+ASC_LEWIS and performing 100 bootstrap replicates. We compared both topologies using Itol v-4.0 viewer (Letunic and Bork 2019).

##### Polymorphisms characterisation

We exported the polymorphic TE positions as an annotation file, and we used bedtools intersect (-wao) to perform their intersection with the reference canonical TE annotation. We then cross-referenced the results with the filtered popoolationTE2 output and defined a decision tree to characterise the TE-polymorphism detected by popoolationTE2 as ‘reference-TE polymorphism’ (ref-polymorphism), ‘extra-detection’, or ‘neo-insertion’ (sup Fig S8).

We considered a reference TE-annotation as polymorphic (e.g. ref-polymorphism locus) if:

i. The position of the polymorphism predicted by PoPoolationTE2 falls between the boundaries of the reference TE-annotation
ii. Both the reference TE-annotation and the predicted polymorphism belong to the same TE-consensus sequence.
iii. The TE has a predicted frequency > 75% in the reference isolate Morelos.

Canonical TE-annotations that did not intersect with polymorphic loci predicted by PopoolationTE2, or that presented frequency variations <1% across the isolates were considered as non-polymorphic.

We classified as ‘neo-insertions’ all the polymorphic loci for which no canonical TE was predicted in the reference annotation (polymorphism position is not included in a reference TE-annotation), but which were detected with a frequency > 25% in at least one isolate different from the reference isolate Morelos, in which the TE frequency should be inferior to 1% and thus considered truly absent in the reference genome.

Finally, we classified as ‘extra-detection’ all the polymorphic loci which did not correspond to a reference annotation but which were detected with a frequency > 25% in the reference isolate Morelos (at least). Polymorphic loci having a frequency between 1% and 25% in Morelos isolate were considered ambiguous and were discarded.

Then, for each TE polymorphism, we investigated the homogeneity of the TE frequency between the isolates.We considered TE frequency was homogeneous between isolates when the maximum frequency variation between isolate was <= to 25%. Above this value, we considered the TE presence frequency was heterogeneous between isolates.

#### Highly Contrasted Polymorphic TE loci (HCPTEs): isolation, characterisation and experimental validation

##### HCPTEs isolation

We considered as highly contrasted all the polymorphic loci for which i) all the isolates had frequency values either < 25% or > 75%, ii) at least one isolate showed a frequency < 25 % while another presented a frequency > 75%. Polymorphic loci fitting with these requirements were exported as an annotation file in the bed format.

##### HCPTEs possible functional impact

We first identified the genes potentially impacted by the HCPTEs by cross-referencing the HCPTEs annotation file with the gene annotation file, using the bedtools suite. We used the ‘closest’ program (-D b - fu -io; b being the gene annotation file) to identify the closest (but not intersecting) gene downstream each HCPTE. We only retained the entries with a maximum distance of 1 kb between the HCPTE and gene boundaries. We identified the insertions in the gene using the ‘intersect’ tool (-wo).

Then, we performed a manual bioinformatic functional analysis for each gene potentially impacted by HCPTEs. Protein sequences were extracted from the *M. incognita* predicted proteome (Blanc-Mathieu et al. 2017) and blasted (blastp; default parameters) against the Non-Redundant protein sequences database (NR) from the NCBI (https://blast.ncbi.nlm.nih.gov/). The same sequences were also used on the InterProScan website (https://www.ebi.ac.uk/interpro/) to perform an extensive search on all the available libraries of conserved protein domains and motifs.

Then, for each gene potentially impacted by HCPTEs, we performed an orthology search on the Wormbase Parasite website (https://parasite.wormbase.org/) using genes accession numbers and the pre-computed ENSEMBL Compara orthology prediction (Herrero et al. 2016).

Finally, we analysed the expression levels of the genes potentially impacted by HCPTEs extracting the information from the RNA-seq analysis of four *M. incognita* life-stages performed previously (see Putative transposition machinery identification section).

##### Experimental validation of Highly Contrasted Polymorphic TE loci

To experimentally validate in-silico predictions of TE neo-insertions with potential functional impact, we selected 5 candidates among the HCPTEs loci and performed a PCR experiment. To run this experiment, we used DNA remaining from extractions performed on the M. incognita isolates for a previous population genomics analysis (Koutsovoulos et al. 2020). We selected loci to be validated based on the following criteria:

- The predicted insertion must be in a genic or potential regulatory region (max 1kb upstream of a gene) as the most evident criterion for a potential functional impact.
- The element must be short enough (2.5kb max) to be amplified by PCR and SANGER sequenced using standard techniques and material.
- To validate the predicted impacted gene actually exists, it must be supported by substantial expression data in the reference isolate Morelos.
- To maximize the chances the genes have effects on biological traits characteristic of the root-knot nematodes, the impacted gene must be *Meloidogyne*-specific.

Once all these criteria were applied, we maximized the diversity of TE orders involved and this resulted in the 5 loci presented in the results section.

#### Primer design and PCR amplification

We designed primers for the PCR analysis using the Primer3Plus web interface (Untergasser et al. 2007). The set of 10 primers with the corresponding sequence and expected amplicon sizes with, or without TE insertion, is shown in (sup. Table 11 & (Kozlowski, Hassanaly-Goulamhoussen, et al. 2020)). We used primers amplifying the whole actin-encoding gene (Minc3s00960g19311) as positive control.

PCR experiments were performed on *M. incognita* Morelos isolate and 11 Brazilian isolates: R1-2, R1-3, R1-6, R2-1, R2-6, R3-1, R3-2, R3-4, R4-1, R4-3 and R4-4.

R3-1 presented no amplification in any of the tested loci nor the positive control (actin) and was thus discarded from this analysis.

PCR mixture contained 0.5µmol of each primer, 1x MyTaq™ reaction buffer and 1.0 U of MyTaq™ DNA polymerase (Bioline Meridian Bioscience) adjusted to a total volume of 20µL. PCR amplification was performed with a TurboCycler2 (Blue-Ray Biotech Corp.). PCR conditions were as follows: initial denaturation at 95°C for 5 min, followed by 35 cycles of 95°C for 30 s, 56°C for 30 s of annealing, and 72°C for 3 min of extension, the program ending with a final extension at 72°C for 10 min. Aliquots of 5µL were migrated by electrophoresis on a 1% agarose gel (Sigma Chemical Co.) for 70 min at 100 V. The size marker used is 1kb Plus DNA Ladder (New England Biolabs Inc.), containing the following size fragments in bp: 100, 200, 300, 400, 500, 600, 700, 900, 1000, 1200, 1500, 2000, 3000, 4000, 5000, 6000, 8000 and 10000.

#### Purification and sequencing of PCR amplicons

Amplicon bands were revealed using ethidium bromide and exposure to ultraviolet radiation. PCR products bands were excised from the agarose gel with a scalpel and purified using MinElute Gel Extraction Kit (Qiagen) before sequencing, following the manufacturer’s protocol. PCR products were sequenced by Sanger Sequencing (Eurofins Genomics).

Forward (F) and Reverse (R) sequences were blasted individually (https://blast.ncbi.nlm.nih.gov/ ; Optimised for ‘Somewhat similar sequences’, default parameters) to the expected TE-consensus sequence and to the genomic region surrounding the predicted insertion point (2 kb region: 1kb upstream the predicted insertion point and 1kb downstream). When no significant hit was found, the sequence was blasted against the *Meloidogyne* reference genomes available (https://meloidogyne.inrae.fr/), the whole TE-consensus library, and the NR database on the NCBI blast website.

## Data accessibility

All the raw and filtered data generated in this study as well as details of the experimental procedures, scripts and codes have been deposited and made publicly available in the institutional INRAE Data Portal at this URL: https://data.inrae.fr/dataverse/TE-mobility-in-MiV3 and cited throughout the text where appropriate, with DOIs available in the references.

## Supplementary material

Supplementary material, tables and figures accompany this article and are available online in bioRxiv as a single PDF file.

## Acknowledgements

The authors would like to thank Joffrey Mejias for all his advice and all the inspiring discussion. The authors are grateful to Laetitia Perfus-Barbeoch for her advice and support in the experimental validation of TE movements. The authors would like to thank Erika VS Albuquerque for her help and assistance in accessing the DNA extractions from the *M. incognita* Brazilian Isolates. We would also like to thank the BIG bioinformatics platform from the PlantBios infrastructure as well as the URGI team for providing facilities and technical support. This work has been supported by the French government, through the UCA-JEDI “Investments in the Future” project managed by the National Research Agency (ANR) with the reference number ANR-15-IDEX-01. Version 4 of this preprint has been peer-reviewed and recommended by Peer Community In Evolutionary Biology (https://doi.org/10.24072/pci.evolbiol.100106)

