## supplementaries for "Transposable Elements are an evolutionary force shaping genomic plasticity in the parthenogenetic root-knot nematode *Meloidogyne incognita*"

* co-last authors
Affiliation : Université Côte d'Azur, INRAE, CNRS, ISA, Sophia Antipolis, France

**Table S1: Per-order summary of *M.incognita* draft TE annotations.**

Autonomous TE orders (*) regroup elements known to present transposition machinery and thus able to transpose by themselves. On the opposite, non-autonomous orders (**) regroup elements lacking transposition machinery and therefore relying on autonomous elements to transpose. "Class 1 & 2 like" regroup elements for which homology-based evidence is sufficient to support an assignment to class I (retro) or II (DNA-transposon), but insufficient to assign a known order. "PotHostGenesOrOther" classification regroups elements which most likely correspond to duplicated genes. "Unclassif." elements are repetitive elements without sufficient evidence to be classified as class I (retro) or II (DNA-transposon). "Class 1 & 2 like", "PotHostGenesOrOther", and "Unclassif." are removed in the canonical TE annotations.

| | **order** autonomous (*) / non-autonomous (**) | **nb. of features** | **total length (bp)** | **genome percentage (%)** | **median length (bp)** | **median identity with consensus (%)** |
|---|---|---|---|---|---|---|
| **Retro - transposon** | SINE (**) | 19 | 6,618 | 0.004 | 258.0 | 87.6 |
| | LARD (**) | 217 | 132,969 | 0.072 | 244.0 | 92.35 |
| | TRIM (**) | 2,466 | 1,240,016 | 0.676 | 468.0 | 76.3 |
| | LINE (*) | 970 | 822,008 | 0.448 | 477.0 | 76.7 |
| | LTR (*) | 2,878 | 2,702,453 | 1.472 | 429.5 | 77.8 |
| **DNA - transposon** | Helitron (*) | 152 | 282,819 | 0.154 | 742.0 | 78.1 |
| | Maverick (*) | 17,684 | 9,553,119 | 5.205 | 364.0 | 74.8 |
| | MITE (**) | 12,435 | 5,126,098 | 2.793 | 363.0 | 88.5 |
| | TIR (**) | 11,094 | 5,389,275 | 2.936 | 379.0 | 85.0 |
| **Others** | CLASS_1_LIKE | 11,053 | 6,737,590 | 3.671 | 522.0 | 74.1 |
| | CLASS_2_LIKE | 77 | 34,339 | 0.019 | 497.0 | 98.7 |
| | potHostGenesOrOther | 26,225 | 12,185,975 | 6.640 | 359.0 | 75.1 |
| | unclassif | 8,811 | 4,212,017 | 2.295 | 390.0 | 79.0 |
| | **Total** | **94,081** | **48,425,296** | **26.385** | | |

**Table S2: Per-order summary of *C.elegans* draft TE annotations.**

| | **order** autonomous (*) / non-autonomous (**) | **nb. of features** | **total length (bp)** | **genome percentage (%)** | **median length (bp)** | **median identity with consensus (%)** |
|---|---|---|---|---|---|---|
| **Retro - transposon** | SINE (**) | 85 | 51,197 | 0.051 | 479.0 | 89.7 |
| | LARD (**) | 14 | 17,043 | 0.017 | 572.5 | 87.8 |
| | TRIM (**) | 3,324 | 2,184,226 | 2.178 | 485.0 | 79.1 |
| | LINE (*) | 519 | 480,089 | 0.479 | 538.0 | 96.4 |
| | LTR (*) | 246 | 215,384 | 0.215 | 509.0 | 96.15 |
| **DNA - transposon** | Helitron (*) | 2,865 | 2,103,981 | 2.098 | 547.0 | 77.2 |
| | Maverick (*) | 26 | 39,843 | 0.040 | 680.0 | 95.25 |
| | MITE (**) | 4,274 | 1,752,665 | 1.748 | 322.0 | 81.4 |
| | TIR (**) | 3,840 | 2,499,195 | 2.492 | 413.0 | 90.45 |
| **Others** | CLASS_1_LIKE | 46 | 19,873 | 0.020 | 385.5 | 85.325 |
| | CLASS_2_LIKE | 5,607 | 1,678,689 | 1.674 | 230.0 | 81.3 |
| | potHostGenesOr Other | 742 | 497,310 | 0.496 | 407.0 | 75.0 |
| | unclassif | 372 | 314,317 | 0.313 | 729.5 | 92.0 |
| | **Total** | **21,960** | **11,853,812** | **11.820** | | |

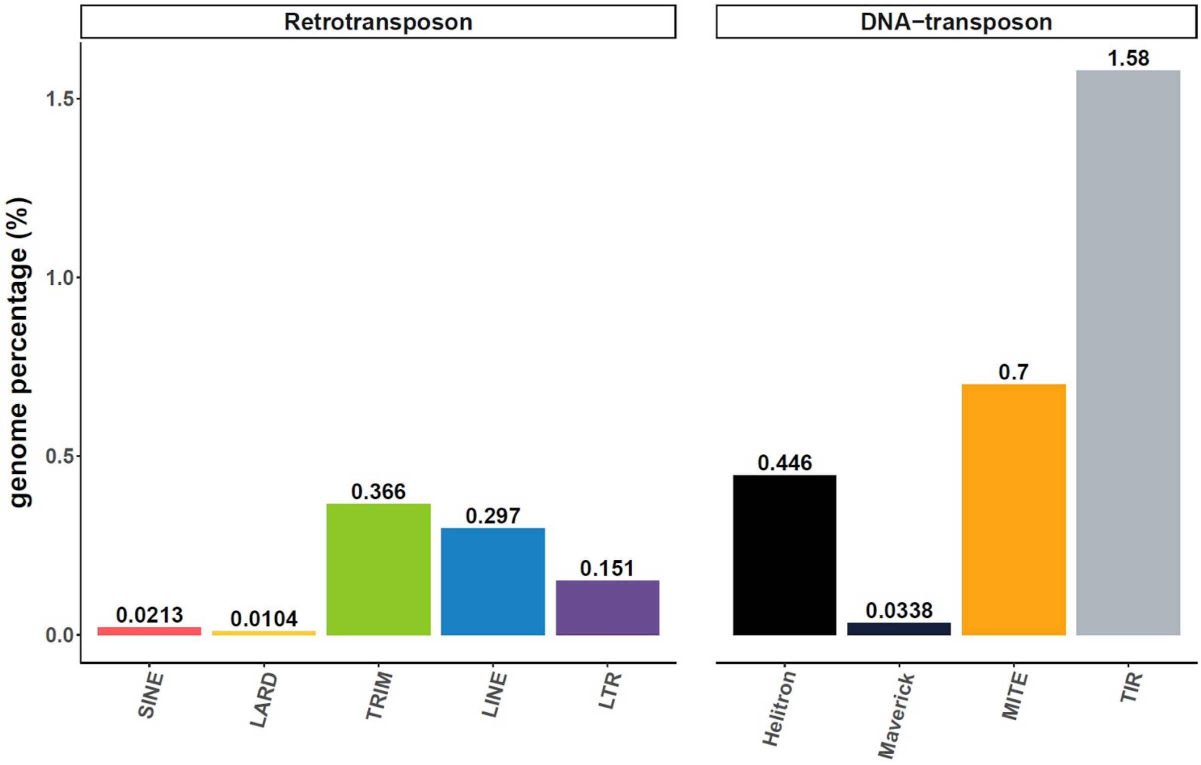


**Fig S1: Canonical TE annotations distribution in the *C. elegans* genome**

Genome percentage is based on a *C. elegans* genome size of 100,286,401 bp.

**Table S3: Per-order summary of *C.elegans* canonical TE annotations.**

| | order<br>autonomous (*)<br>/<br>non-autonomous (**) | nb. of features | total length (bp) | genome percentage (%) | median length (bp) | median identity with consensus (%) |
|---|---|---|---|---|---|---|
| **Retro - transposon** | SINE (**) | 23 | 21,342 | 0.021 | 908.0 | 98.1 |
| | LARD (**) | 3 | 10,417 | 0.010 | 3969.0 | 99.7 |
| | TRIM (**) | 294 | 366,742 | 0.366 | 744.5 | 90.6 |
| | LINE (*) | 184 | 297,840 | 0.297 | 1252.5 | 98.7 |
| | LTR (*) | 124 | 151,145 | 0.151 | 617.5 | 97.75 |
| **DNA - transposon** | Helitron (*) | 267 | 447,385 | 0.446 | 1514.0 | 96.1 |
| | Maverick (*) | 14 | 33,884 | 0.034 | 1399.5 | 97.6 |
| | MITE (**) | 1,101 | 702,012 | 0.700 | 521.0 | 95.0 |
| | TIR (**) | 1,475 | 1,582,321 | 1.578 | 815.0 | 97.1 |
| | **Total** | **3,485** | **3,613,088** | **3.603** | | |

**Table S4:** ***M. incognita*** **per-order summary of copies % identity with their consensus.**

| | Min. | 1st Quantile | Median | Mean | 3rd Quantile | Max. |
|---|---|---|---|---|---|---|
| Helitron | 85.3 | 92.0 | 94.4 | 93.4 | 95.8 | 97.7 |
| LARD | 92.6 | 96.1 | 97.1 | 96.9 | 97.9 | 99 |
| LINE | 85.9 | 95.4 | 96.6 | 96.3 | 98.8 | 100 |
| LTR | 85.3 | 94.8 | 97.0 | 96.3 | 98.4 | 100 |
| Maverick | 85.0 | 90.3 | 95.3 | 93.7 | 97.2 | 99.8 |
| MITE | 85.0 | 92.8 | 96.2 | 95.3 | 98.4 | 100 |
| SINE | 93.4 | 99.3 | 99.7 | 98.9 | 99.8 | 100 |
| TIR | 85.0 | 93.7 | 97.3 | 96.0 | 99.2 | 100 |
| TRIM | 85.2 | 95.7 | 97.7 | 96.8 | 98.8 | 99.9 |

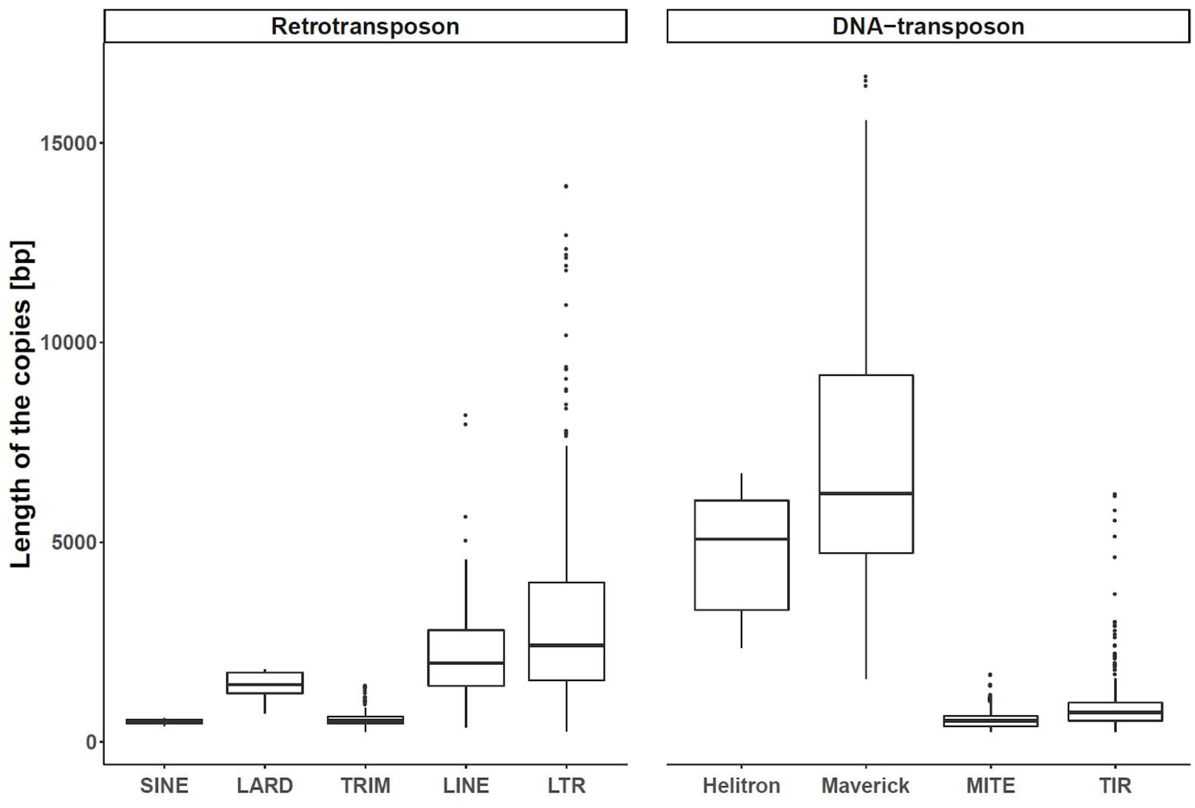


**Fig S2: Distribution of TE lengths per-order (*M. incognita*).**

Box plots per order of the distribution of the canonical TE annotations' length .

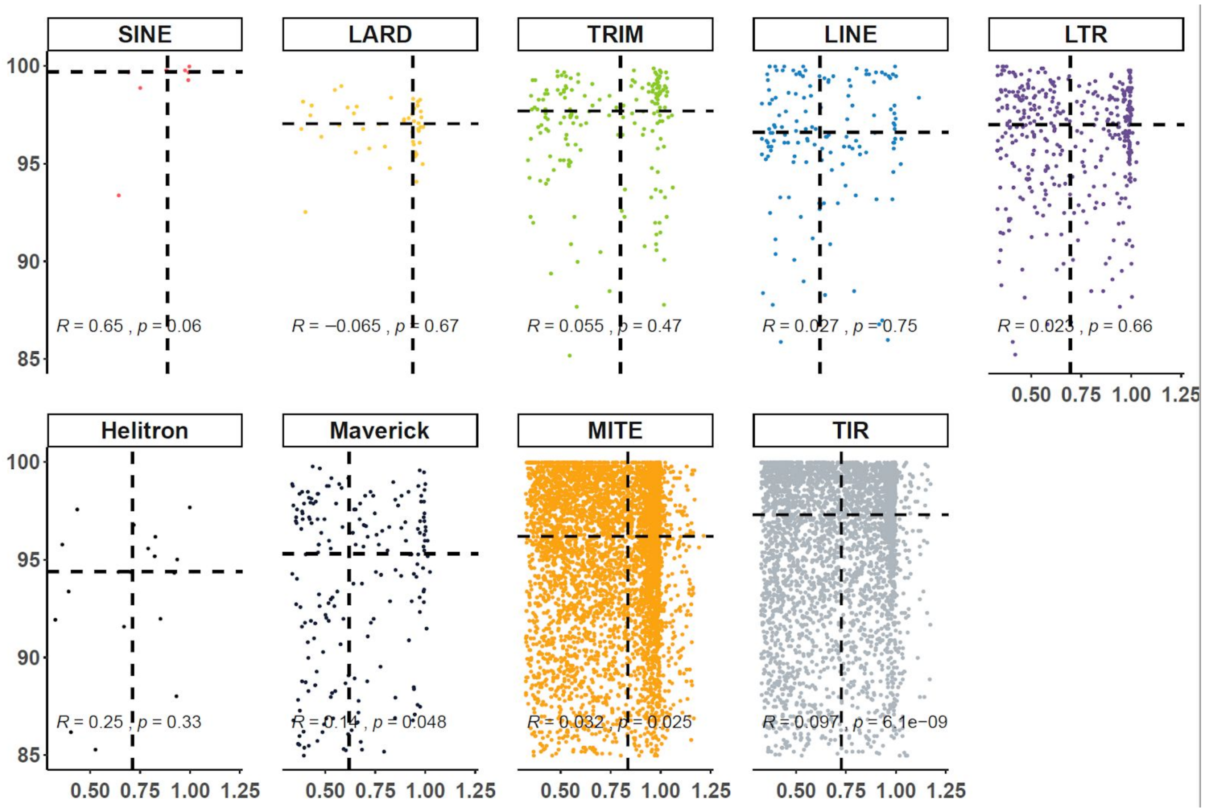


**Fig S3: *M. incognita* per TE copy % identity with its consensus in function of the proportion of consensus covered.**

Data are splitted in panels according to TE orders. For each panel, Y-axis represents the percentage of identity a copy shares with its consensus. X-axis represents the coverage of the TE consensus (proportion). Coverage values > 100% correspond to cases for which the copy includes a nested sequence regarding the TE consensus sequence (other TE, repeats, other). Each point represents a TE locus (*i.e.* a TE copy). Dashed lines represent the per order median value of both the identity percentage (horizontal line) and the proportion of coverage (vertical line). R value represents the correlation coefficient (Pearson) computed for each order, and p is the associated p-value.

**Table S5: canonical TE annotations with putative transposition machinery**

| | autonomous (*) / non-autonomous (**) orders | nb. of annotations with putative transposition machinery | nb. of annotations with substantially expressed putative transposition machinery |
|---|---|---|---|
| retro - transposon | SINE (**) | 0 | 0 |
| | LARD (**) | 0 | 0 |
| | TRIM (**) | 0 | 0 |
| | LINE (*) | 54 | 26 |
| | LTR (*) | 147 | 45 |
| DNA - transposon | Helitron (*) | 17 | 3 |
| | Maverick (*) | 63 | 26 |
| | MITE (**) | 0 | 0 |
| | TIR (*) | 30 | 6 |

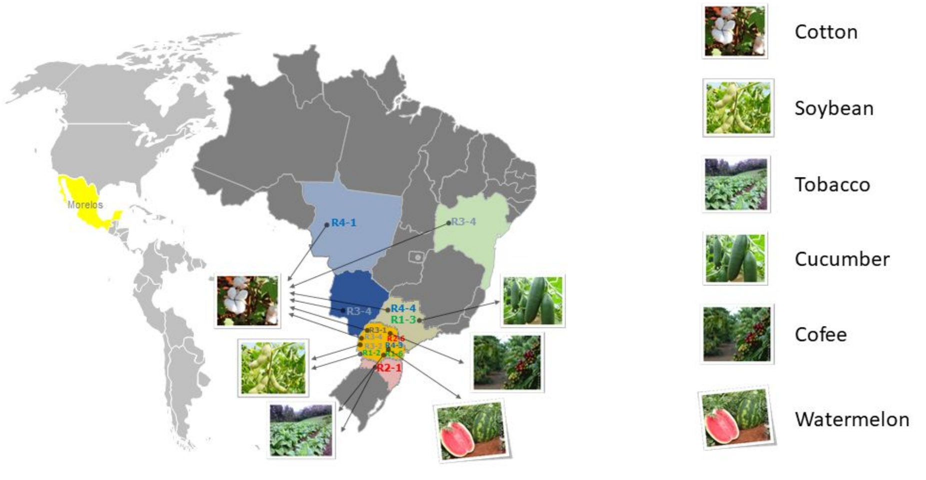


Adapted from Evolutionary Applications, Volume: 13, Issue: 2, Pages: 442-457, First published: 19 October 2019, DOI: (10.1111/eva.12881)

**Fig S4: Isolates geographical distribution and host plants.**

American continent map showing the geographical distribution for all isolates used in the study. Expanded map of Brazil shows the states where the 11 isolates sequenced in (Koutsovoulos et al. 2020) were collected. Each state is highlighted with a different colour. The crops from which the samples were isolated are illustrated by photographs, which are pointed by arrows coming from the name of the respective isolate.

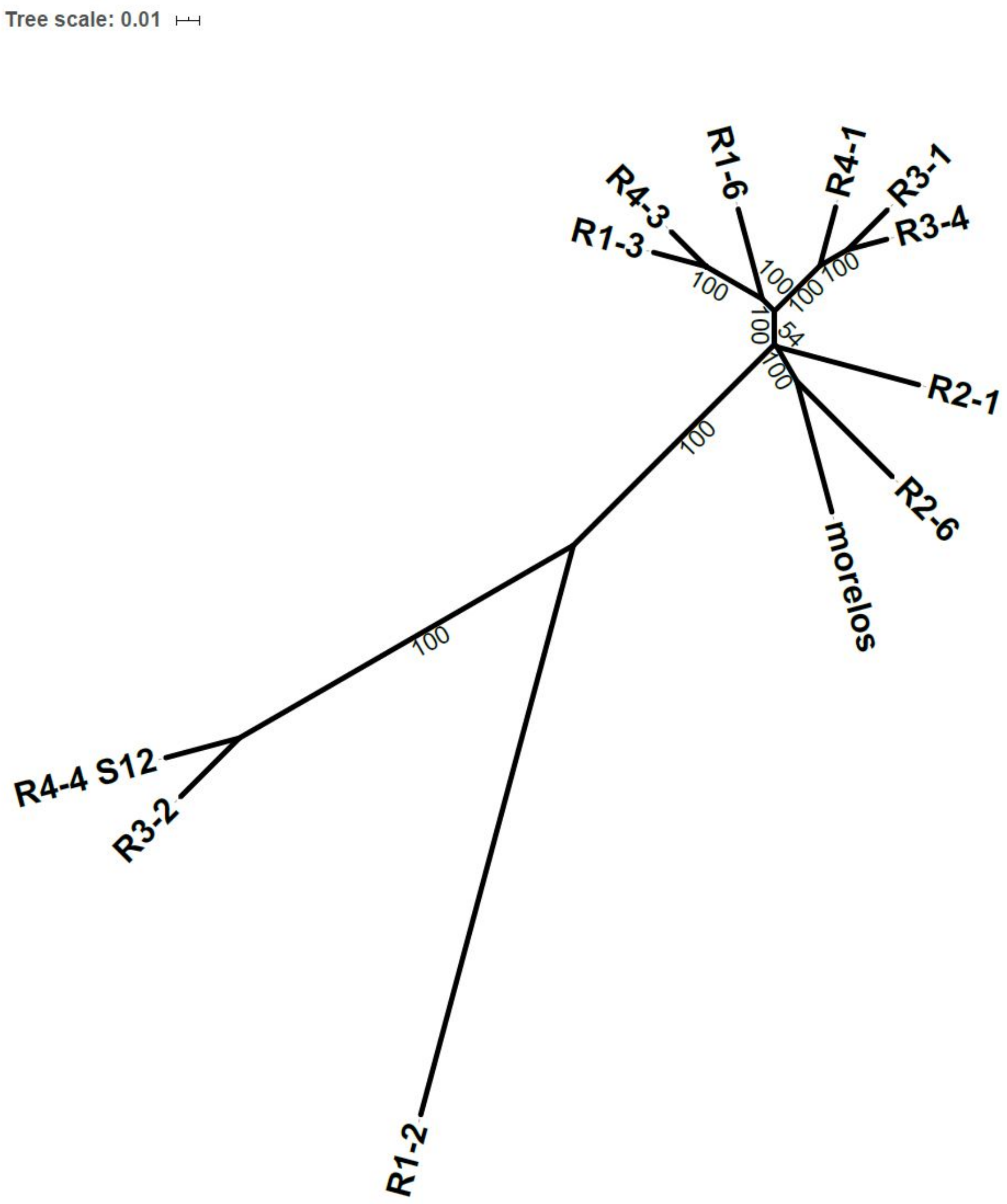
Tree scale: 0.01
R4-3
R1-6
R4-1
R3-1
R3-4
R1-3
100
54
R2-1
R2-6
morelos
R4-4 S12
R3-2
R1-2

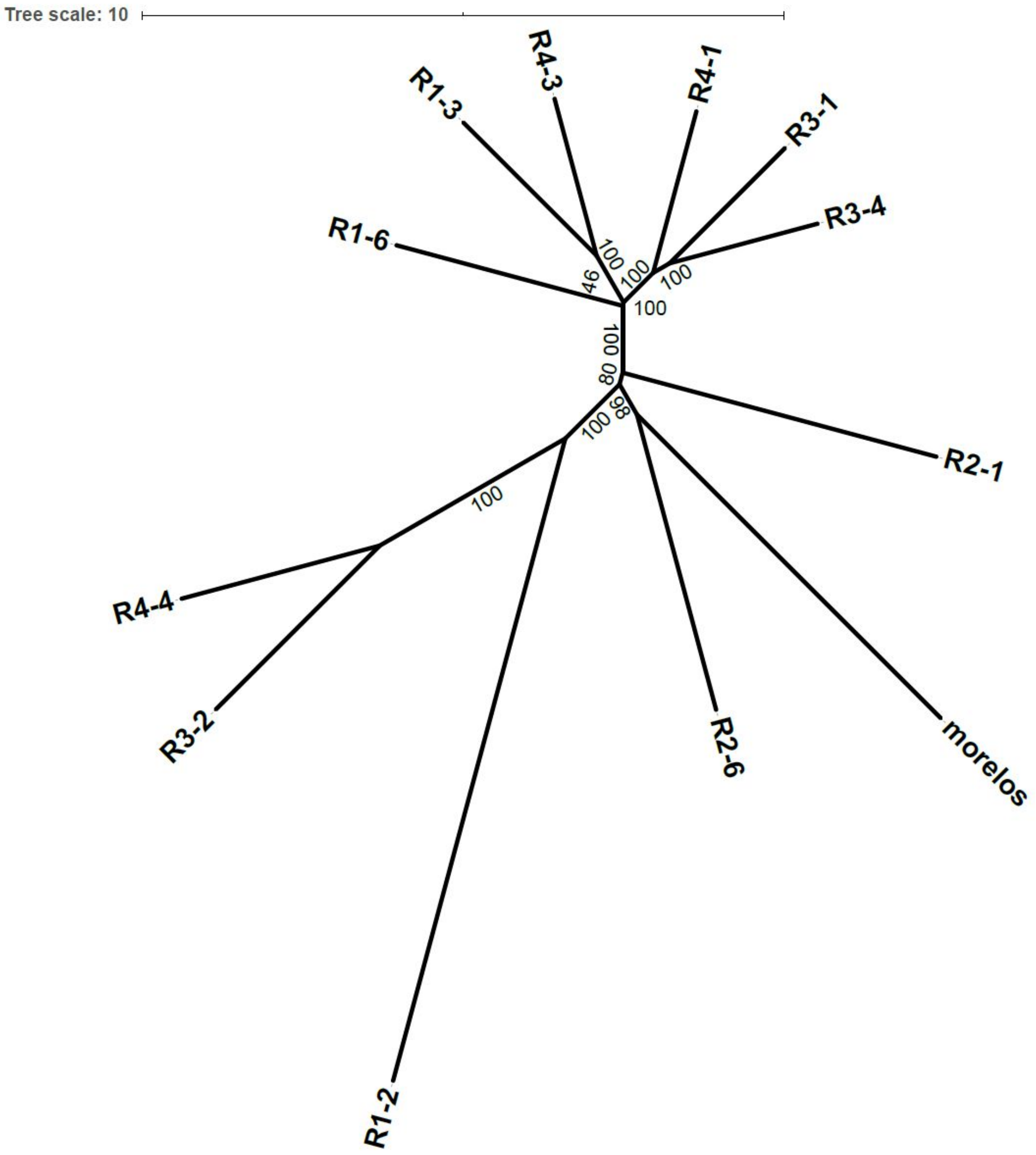


**Fig S5: Phylogenetic tree for *M. incognita* isolates with branch length.**

Top Phylogenetic tree is based on SNP present in coding sequences. Maximum Likelihood (ML) tree reconstruction. Bottom Phylogenetic tree based on TE-frequencies euclidean distances between isolates. Neighbor-Joining (NJ) tree reconstruction. For both trees, the branch length is displayed. The scale is displayed in the top-left corner of each sub-figure. Values in black (bootstrap values) represent nodes support values, 100 being the maximal value.

**Table S6: TE repartition per orders in the reference annotation and between polymorphisms types.**

Ref-annotation line represents the per-order number of elements in the reference genome annotation. The sum of "non-polymorphic ref." and "polymorphic-ref" is not equal to the number of reference annotations due to filtering criteria. See sup. Fig S8 for detailed explanations.

| | SINE | LARD | TRIM | LINE | LTR | Helitron | Maverick | MITE | TIR |
|---|---|---|---|---|---|---|---|---|---|
| **ref-annotations** | 9 | 45 | 174 | 145 | 373 | 18 | 189 | 5085 | 3595 |
| **non-polymorphic ref-annotations** | 8 | 35 | 154 | 128 | 322 | 16 | 179 | 3602 | 2657 |
| **polymorphic ref-annotations** | 1 | 6 | 14 | 13 | 37 | 1 | 7 | 1194 | 818 |
| **neo-insertions** | 0 | 0 | 0 | 10 | 11 | 0 | 0 | 192 | 74 |
| **extra-detection** | 0 | 0 | 4 | 6 | 12 | 2 | 16 | 97 | 69 |

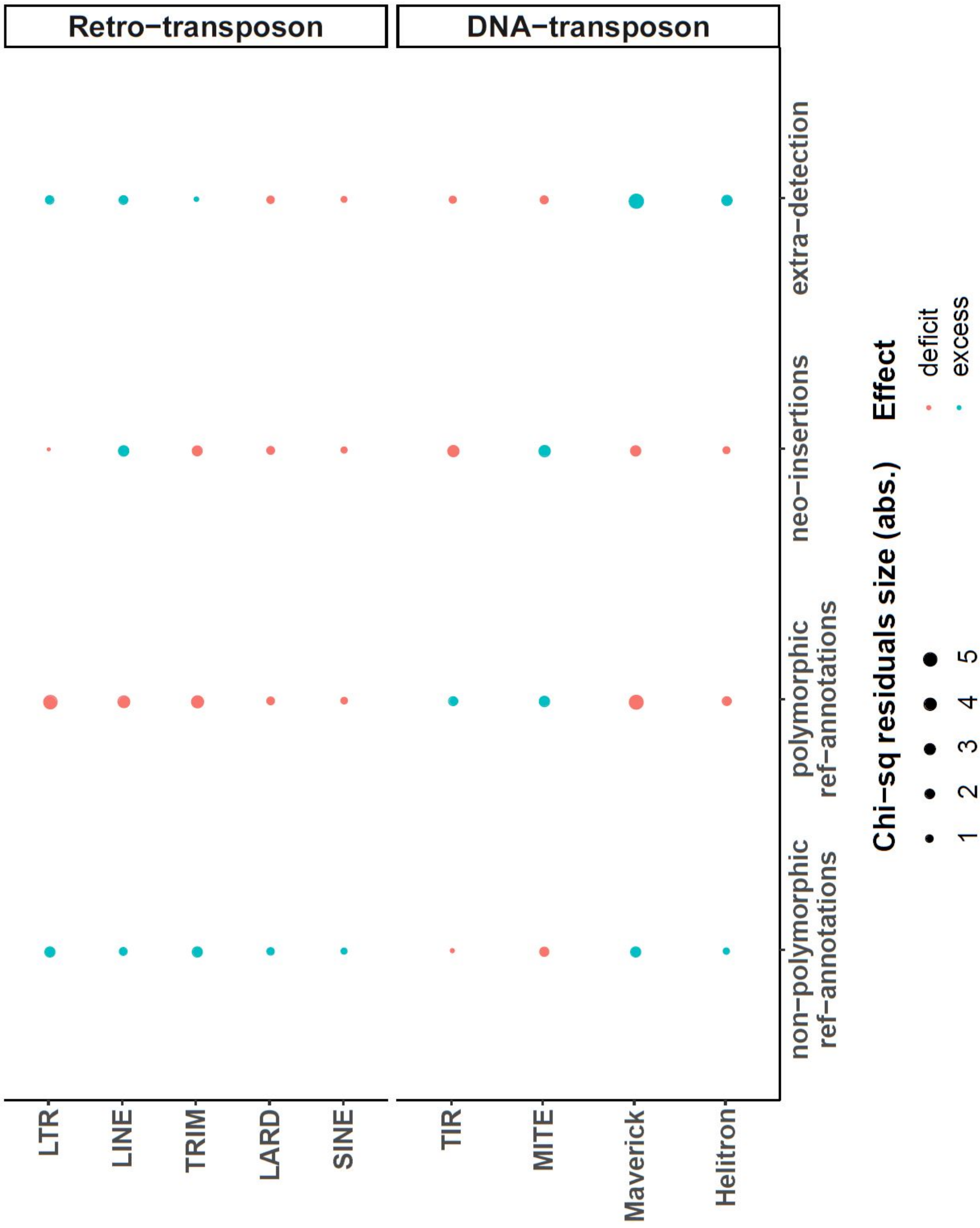


**Fig S6: Relative abundance of the TE copies (count) per polymorphism types and TE-orders.**

Each point represents a chi-square residual value. Chi-square residuals are the distance from the expected distribution under the homogeneity hypothesis. They are used here as a proxy to estimate the relative abundance (count) per polymorphism type and TE order and how each combination differs from the expected distribution. For each point, the wider is the surface, the higher is the distance from the expectation. Red points represent a deficit compared to the expectation while the blue points represent an excess.

**Table S7: Number of HCPTEs copies per-consensus.**

| consensus | order | nb. of HCPTEs copies |
|---|---:|---:|
| DTX-comp_mincV3XDN-B-R1459-Map20 | TIR | 8 |
| DTX-incomp_mincV3XDN-B-R11531-Map10 | TIR | 2 |
| DTX-incomp_mincV3XDN-B-R271-Map10 | TIR | 1 |
| DTX-incomp_mincV3XDN-B-R3892-Map20 | TIR | 1 |
| DXX-MITE_mincV3XDN-B-G1048-Map15 | MITE | 1 |
| DXX-MITE_mincV3XDN-B-G305-Map9 | MITE | 1 |
| DXX-MITE_mincV3XDN-B-R14125-Map7 | MITE | 1 |
| DXX-MITE_mincV3XDN-B-R306-Map20 | MITE | 10 |
| DXX-MITE_mincV3XDN-B-R321-Map20 | MITE | 1 |
| DXX-MITE_mincV3XDN-B-R3266-Map20 | MITE | 1 |
| DXX-MITE_mincV3XDN-B-R3611-Map9 | MITE | 4 |
| RIX-comp_mincV3XDN-B-R6875-Map20_reversed | LINE | 1 |
| RIX-incomp_mincV3XDN-B-R4613-Map9 | LINE | 1 |

**Table S8: Orthologs to genes potentially impacted by HCPTEs.**

Entries with bold font are *M. incognita*'s genes with orthologs in other *Meloidogyne species* or other Plant Parasitic Nematode (PPN) genus only. Entries are sorted by gene name. Genes highlighted in yellow are the genes potentially impacted by HCPTEs which have been selected for experimental validation.

| Gene | Wormbase gene trees orthologs | Which species | Tree URL |
|---|---|---|---|
| Minc3s00005g00347 | 154 | in many nematodes and other animals | https://parasite.wormbase.org/Meloidogyne_incognita_prjeb8714/Gene/Compara_Tree?g=Minc3s00005g00347;r=FXSY01000005.1:267231-279946;t=Minc3s00005g00347 |
| Minc3s00005g00348 | 168 | in many nematodes and other animals | https://parasite.wormbase.org/Meloidogyne_incognita_prjeb8714/Gene/Compara_Tree?g=Minc3s00005g00348;r=FXSY01000005.1:271509-282281;t=Minc3s00005g00348 |
| **Minc3s00026g01668** | 14 | **Meloidogyne-specific: incognita, arenaria, javanica, floridensis** | https://parasite.wormbase.org/Meloidogyne_incognita_prjeb8714/Gene/Compara_Tree?g=Minc3s00026g01668;r=FXSY01000026.1:125149-126932;t=Minc3s00026g01668;collapse= |
| Minc3s00137g05752 | 122 | in many nematodes and other animals | https://parasite.wormbase.org/Meloidogyne_incognita_prjeb8714/Gene/Compara_Tree?g=Minc3s00137g05752;r=FXSY01000137.1:70079-73404;t=Minc3s00137g05752 |
| Minc3s00157g06330 | 135 | in many nematodes and other animals | https://parasite.wormbase.org/Meloidogyne_incognita_prjeb8714/Gene/Compara_Tree?g=Minc3s00157g06330;r=FXSY01000157.1:83470-88312;t=Minc3s00157g06330 |
| Minc3s00199g07364 | 203 | in many nematodes and other animals | https://parasite.wormbase.org/Meloidogyne_incognita_prjeb8714/Gene/Compara_Tree?g=Minc3s00199g07364;r=FXSY01000199.1:14729-17937;t=Minc3s00199g07364 |
| Minc3s00199g07365 | 149 | nematode specific but many nematodes | https://parasite.wormbase.org/Meloidogyne_incognita_prjeb8714/Gene/Compara_Tree?g=Minc3s00199g07365;r=FXSY01000199.1:18300-23780;t=Minc3s00199g07365 |
| Minc3s00201g07425 | 188 | in many nematodes and other animals | https://parasite.wormbase.org/Meloidogyne_incognita_prjeb8714/Gene/Compara_Tree?g=Minc3s00201g07425;r=FXSY01000201.1:30179-31671;t=Minc3s00201g07425 |
| Minc3s00201g07426 | | tRNA (non-coding), widely conserved in nematodes | |
| Minc3s00201g07427 | 5 | nematode specific, mainly Meloidogyne but also Chromadorea and Dirofilaria | https://parasite.wormbase.org/Meloidogyne_incognita_prjeb8714/Gene/Compara_Tree?g=Minc3s00201g07427;r=FXSY01000201.1:31822-32328;t=Minc3s00201g07427 |

| | | | |
|---|---|---|---|
| Minc3s00301g09724 | 129 | in many nematodes and other animals | https://parasite.wormbase.org/Meloidogyne_incognita_prjeb8714/Gene/Compara_Tree?g=Minc3s00301g09724;r=FXSY01000301.1:27845-35780;t=Minc3s00301g09724 |
| **Minc3s00450g12515** | 5 | **Meloidogyne-specific: incognita, arenaria.** | https://parasite.wormbase.org/Meloidogyne_incognita_prjeb8714/Gene/Compara_Tree?g=Minc3s00450g12515;r=FXSY01000450.1:51949-52954;t=Minc3s00450g12515;collapse= |
| **Minc3s00621g15225** | 9 | **Meloidogyne-specific: incognita, arenaria, javanica, floridensis, enterolobii, hapla, graminicola.** | https://parasite.wormbase.org/Meloidogyne_incognita_prjeb8714/Gene/Compara_Tree?g=Minc3s00621g15225;r=FXSY01000621.1:38374-38735;t=Minc3s00621g15225 |
| Minc3s00667g15847 | 17 | nematode specific, all Plant Parasitic Nematodes (PPN) except A. nanus | https://parasite.wormbase.org/Meloidogyne_incognita_prjeb8714/Gene/Compara_Tree?g=Minc3s00667g15847;r=FXSY01000667.1:10892-13619;t=Minc3s00667g15847 |
| **Minc3s00751g16867** | 13 | **Meloidogyne-specific: incognita, arenaria, javanica, floridensis, enterolobii.** | https://parasite.wormbase.org/Meloidogyne_incognita_prjeb8714/Gene/Compara_Tree?g=Minc3s00751g16867;r=FXSY01000751.1:15531-16499;t=Minc3s00751g16867 |
| Minc3s00905g18730 | 251 | in many nematodes and other animals | https://parasite.wormbase.org/Meloidogyne_incognita_prjeb8714/Gene/Compara_Tree?g=Minc3s00905g18730;r=FXSY01000905.1:1630-5121;t=Minc3s00905g18730 |
| **Minc3s00905g18731** | 5 | **Meloidogyne-specific: incognita, arenaria, javanica, floridensis, enterolobii.** | https://parasite.wormbase.org/Meloidogyne_incognita_prjeb8714/Gene/Compara_Tree?g=Minc3s00905g18731;r=FXSY01000905.1:5454-6420;t=Minc3s00905g18731 |
| **Minc3s00909g18773** | 14 | **Meloidogyne-specific: incognita, arenaria, javanica, floridensis, enterolobii, graminicola.** | https://parasite.wormbase.org/Meloidogyne_incognita_prjeb8714/Gene/Compara_Tree?g=Minc3s00909g18773;r=FXSY01000909.1:23309-24625;t=Minc3s00909g18773 |
| Minc3s00965g19365 | 160 | in many nematodes and other animals | https://parasite.wormbase.org/Meloidogyne_incognita_prjeb8714/Gene/Compara_Tree?g=Minc3s00965g19365;r=FXSY01000965.1:6357-15984;t=Minc3s00965g19365 |
| **Minc3s00988g19605** | 3 | **Meloidogyne-specific: incognita, arenaria, javanica.** | https://parasite.wormbase.org/Meloidogyne_incognita_prjeb8714/Gene/Compara_Tree?g=Minc3s00988g19605;r=FXSY01000988.1:15653-17968;t=Minc3s00988g19605 |
| **Minc3s01127g20975** | 4 | **Meloidogyne-specific: incognita, arenaria, javanica, floridensis, hapla.** | https://parasite.wormbase.org/Meloidogyne_incognita_prjeb8714/Gene/Compara_Tree?g=Minc3s01127g20975;r=FXSY01001127.1:7481-15500;t=Minc3s01127g20975 |
| **Minc3s01138g21099** | 6 | **Meloidogyne-specific: incognita, arenaria, javanica, floridensis, enterolobii, hapla, graminicola.** | https://parasite.wormbase.org/Meloidogyne_incognita_prjeb8714/Gene/Compara_Tree?g=Minc3s01138g21099;r=FXSY01001138.1:37292-39709;t=Minc3s01138g21099 |

| | | | |
|---|---|---|---|
| Minc3s01318g22714 | 10 | **PPN-specific: i) Meloidogyne: incognita, arenaria, javanica, floridensis, enterolobii, graminicola; ii) Globobodera: rostochiensis; iii) Ditylenchus: destructor** | https://parasite.wormbase.org/Meloidogyne_incognita_prjeb8714/Gene/Compara_Tree?g=Minc3s01318g22714;r=FXSY01001318.1:1931-3523;t=Minc3s01318g22714 |
| Minc3s01455g23950 | 0 | no gene tree at all: gene specific to Meloidogyne incognita | |
| **Minc3s01827g26567** | 3 | **Meloidogyne-specific: incognita, arenaria, javanica, floridensis, enterolobii.** | https://parasite.wormbase.org/Meloidogyne_incognita_prjeb8714/Gene/Compara_Tree?g=Minc3s01827g26567;r=FXSY01001827.1:2587-2859;t=Minc3s01827g26567;collapse=9293401 |
| Minc3s02496g30324 | 170 | in many nematodes and other animals | https://parasite.wormbase.org/Meloidogyne_incognita_prjeb8714/Gene/Compara_Tree?g=Minc3s02496g30324;r=FXSY01002496.1:10099-17217;t=Minc3s02496g30324 |
| Minc3s03567g34213 | 78 | Present in many animals then only Meloidogyne | https://parasite.wormbase.org/Meloidogyne_incognita_prjeb8714/Gene/Compara_Tree?g=Minc3s03567g34213;r=FXSY01003567.1:9085-10645;t=Minc3s03567g34213;collapse=14989368.14989316.14989313.14989310.14988329 |

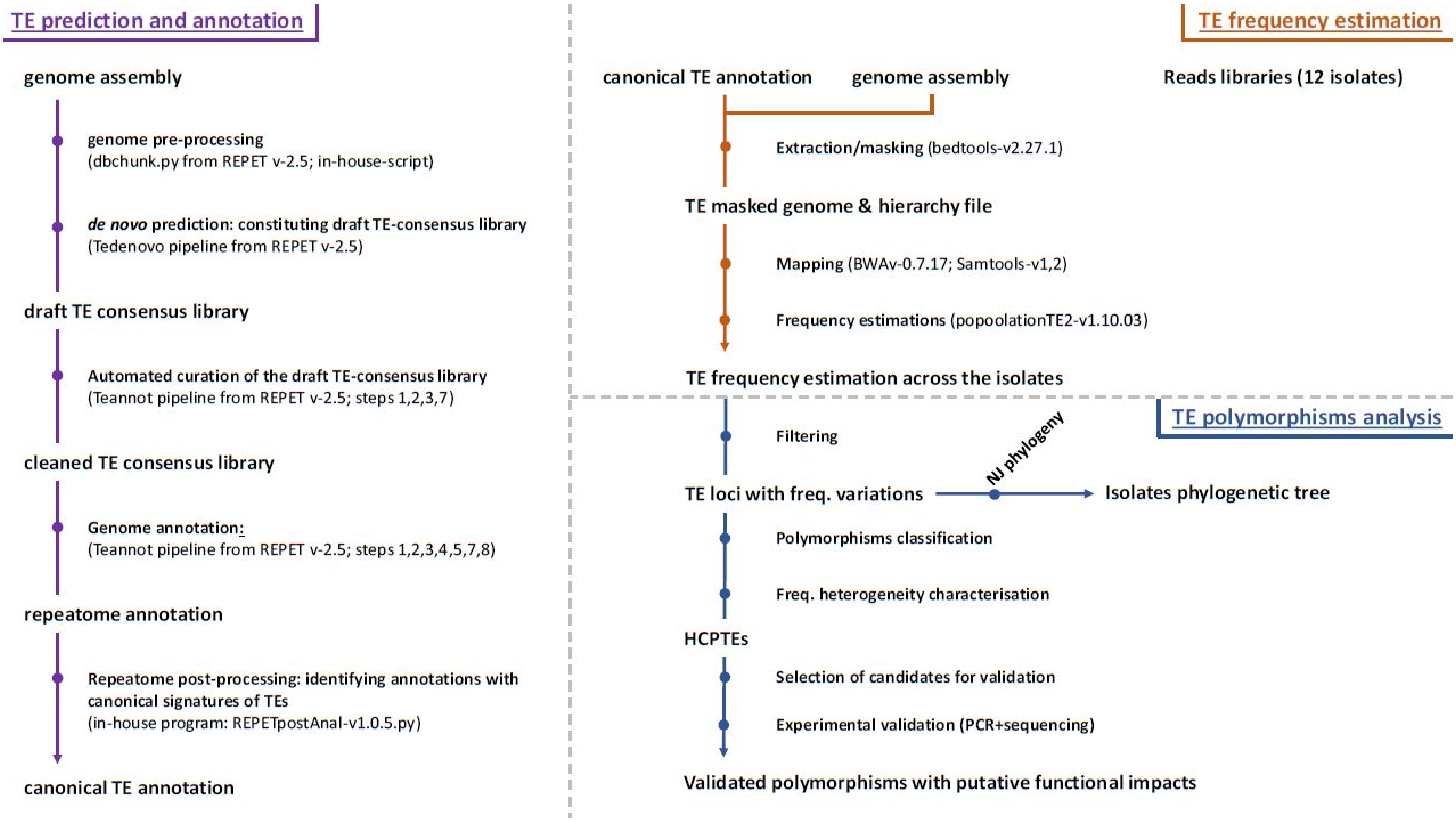
TE prediction and annotation
genome assembly
genome pre-processing
(dbchunk.py from REPET v-2.5; in-house-script)
de novo prediction: constituting draft TE-consensus library
(Tedenovo pipeline from REPET v-2.5)
draft TE consensus library
Automated curation of the draft TE-consensus library
(Teannot pipeline from REPET v-2.5; steps 1,2,3,7)
cleaned TE consensus library
Genome annotation:
(Teannot pipeline from REPET v-2.5; steps 1,2,3,4,5,7,8)
repeatome annotation
Repeatome post-processing: identifying annotations with
canonical signatures of TEs
(in-house program: REPETpostAnal-v1.0.5.py)
canonical TE annotation
TE frequency estimation
canonical TE annotation
genome assembly
Reads libraries (12 isolates)
Extraction/masking (bedtools-v2.27.1)
TE masked genome & hierarchy file
Mapping (BWAv-0.7.17; Samtools-v1,2)
Frequency estimations (popoolationTE2-v1.10.03)
TE frequency estimation across the isolates
TE polymorphisms analysis
Filtering
TE loci with freq. variations
NJ phylogeny
Isolates phylogenetic tree
Polymorphisms classification
Freq. heterogeneity characterisation
HCPTEs
Selection of candidates for validation
Experimental validation (PCR+sequencing)
Validated polymorphisms with putative functional impacts

**Fig S7: workflow overview.**

The current analysis encompasses 3 pipelines: the TE prediction and annotation, the TE frequency estimation, and the TE polymorphisms analysis. Each step's workflow is represented in a separated panel. Each step of each sub-pipeline is explained in detail in Methods. All the scripts are available in (Kozlowski 2020). "Polymorphisms classification" and "Freq. heterogeneity characterisation" steps of the TE-polymorphism pipeline are detailed as a decision tree in Fig S8

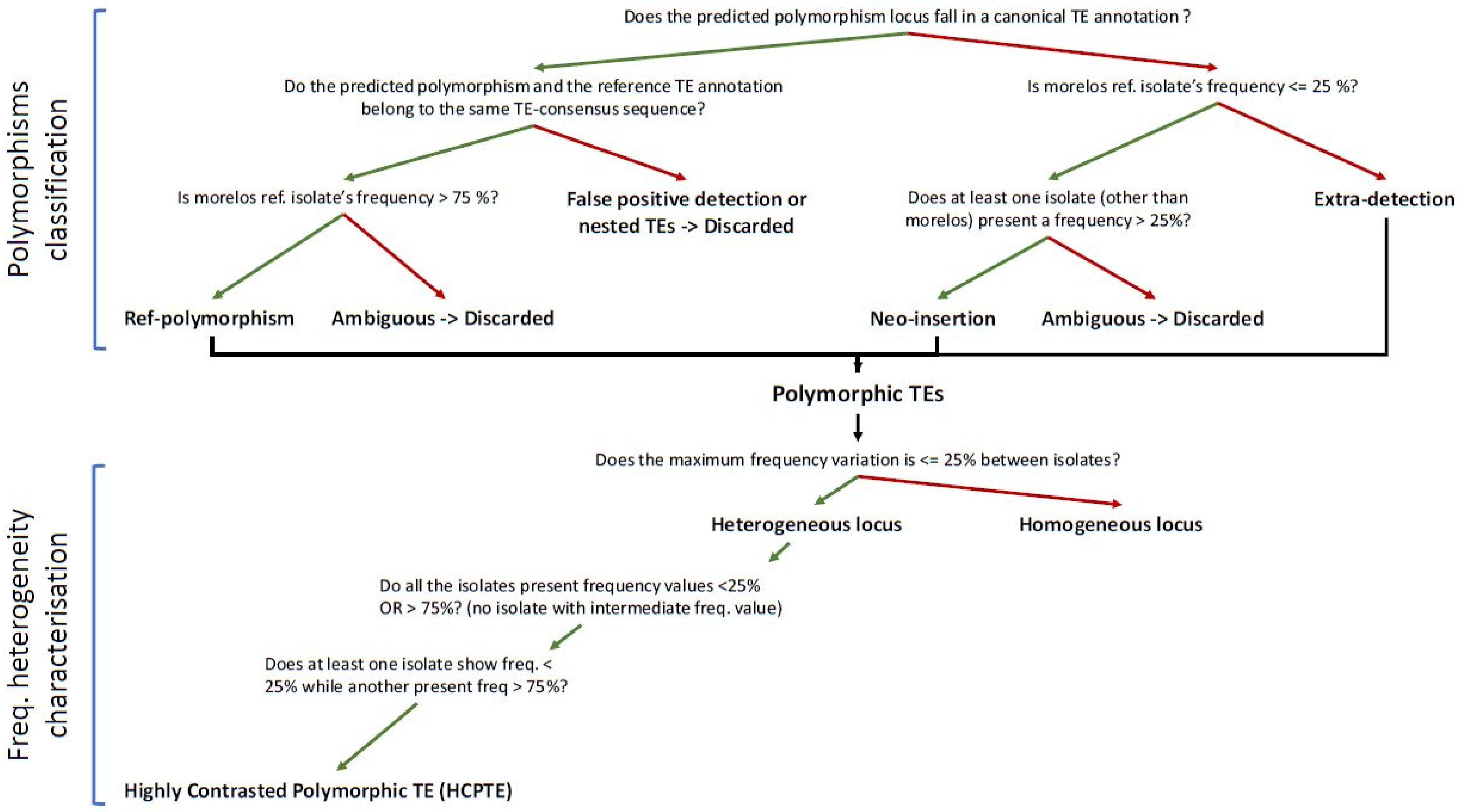

Polymorphisms classification
Does the predicted polymorphism locus fall in a canonical TE annotation ?
Do the predicted polymorphism and the reference TE annotation belong to the same TE-consensus sequence?
Is morelos ref. isolate's frequency <= 25 %?
Is morelos ref. isolate's frequency > 75 %?
False positive detection or nested TEs -> Discarded
Does at least one isolate (other than morelos) present a frequency > 25%?
Extra-detection
Ref-polymorphism
Ambiguous -> Discarded
Neo-insertion
Ambiguous -> Discarded
Polymorphic TEs
Freq. heterogeneity characterisation
Does the maximum frequency variation is <= 25% between isolates?
Heterogeneous locus
Homogeneous locus
Do all the isolates present frequency values <25% OR > 75%? (no isolate with intermediate freq. value)
Does at least one isolate show freq. < 25% while another present freq > 75%?
Highly Contrasted Polymorphic TE (HCPTE)

**FIG S8: decision trees for polymorphisms classification and frequency heterogeneity characterisation.**

This figure details as a decision tree the "Polymorphisms classification" and "Freq. heterogeneity characterisation" steps of the TE-polymorphism pipeline from the Fig S7. Green arrows represent a positive answer. The red ones represent a negative answer.

**Table S9: pairwise blastn of locus 1 sequencing results.**

| insertion predicted | subject | query | sequence | %identity | query cover (%) | e-value | average % identity |
|---|---|---|---|---|---|---|---|
| N | morelos | R2-1 | F | 98.68 | 99 | 6,00E-75 | 99.34 |
| | | | R | 100 | 91 | 2,00E-73 | |
| N | morelos | R2-6 | F | 100 | 93 | 8,00E-73 | 95.3 |
| | | | R | 90.6 | 89 | 4,00E-52 | |
| N | R2-6 | R2-1 | F | 98.64 | 96 | 1,00E-71 | 94.45 |
| | | | R | 90.26 | 92 | 4,00E-52 | |
| Y | R1-2 | R4-4 | F | 92.73 | 72 | 0,00E+00 | 96.25 |
| | | | R | 99.77 | 99 | 0,00E+00 | |
| Y | R3-2 | R1-2 | F | 98.52 | 98 | 0,00E+00 | 98.88 |
| | | | R | 99.24 | 89 | 0,00E+00 | |
| Y | R4-4 | R3-2 | F | 92.16 | 93 | 0,00E+00 | 95.7 |
| | | | R | 99.24 | 98 | 0,00E+00 | |

**Table S10: Reads libraries accession numbers & statistics**

| Lib. name | acess. nb. (SRA) | nb. reads (P-E) | read length (bp) | % GC |
|---|---|---|---|---|
| morelos | ERS1696677 | 76077411 | 2*150 | 28 |
| R1-2 | SRX4373671 | 76359269 | 2*150 | 29 |
| R1-3 | SRX4373672 | 75542522 | 2*150 | 28 |
| R1-6 | SRX4373673 | 75033425 | 2*150 | 28 |
| R2-1 | SRX4373674 | 75065658 | 2*150 | 29 |
| R2-6 | SRX4373675 | 75300726 | 2*150 | 29 |
| R3-1 | SRX4373676 | 74468408 | 2*150 | 30 |
| R3-2 | SRX4373677 | 74671928 | 2*150 | 28 |
| R3-4 | SRX4373678 | 74620706 | 2*150 | 28 |
| R4-1 | SRX4373679 | 75063890 | 2*150 | 28 |
| R4-3 | SRX4373680 | 75235737 | 2*150 | 29 |
| R4-4 | SRX4373681 | 74987959 | 2*150 | 28 |

**Table S11: PCR primers targeting 5 candidate locus for TE insertion**

| Primer Pair | Sequence | Amplicon size without insertion (bp) | Amplicon size with insertion (bp) |
|---|---|---|---|
| Locus1-F | CTTAGGTTTTTGACTGCGTCTGCCAT | 180 | 973 |
| Locus1-R | CAGATGCATTGCGGTGACGTTCTT | | |
| Locus2-F | GGGGGTCAGATTACCCTCTATTATGGCA | 761 | 1870 |
| Locus2-R | CCTCTCCCATCACTCTCACAACCCA | | |
| Locus3-F | CCGTCGGCGGGATCCCTGATATAAA | 690 | 1814 |
| Locus3-R | TTATCGGTTTCAACCCCGACCGAAC | | |
| Locus4-F | GGTGGTGTTGTTGCTGGAATTACTAACC | 981 | 1781 |
| Locus4-R | GACAAACGTTGGAGCACGTTATGCTCG | | |
| Locus5-F | GGAACAGTCAGCGGTGTCGGAAATC | 1005 | 2080 |
| Locus5-R | GTGTATGCTTCAGAACCCAGACGGGGA | | |
| actin-F (ctrl +) | AAGATGGATGAAGAGGTAGCCGCCC | - | 1667 |
| actin-R (ctrl +) | ACTCTTGCTTGCTGATCCACCTTGA | | |
